# Loss of tight junction protein claudin 18 uncovers alveolar epithelial stem cell plasticity and emergence of non-fibrogenic transitional progenitors

**DOI:** 10.1101/2025.10.01.679859

**Authors:** A Castaldi, JS Chin, B Ma, MW Chang, K Samimi, A Allen, W Pinson-Rose, J Castillo, P Zamani, M Xie, N Arias, K Solaimanpour, Y Liu, H Shen, BT Scott, M Horie, P Flodby, A Kasirer-Friede, K Dang, J Buchanan, R Lancione, Michael Miller, Qian Yang, C Benner, C Marconett, A Wang, B Zhou, X Sun, Z Borok

## Abstract

Persistence of senescent alveolar transitional progenitors following lung injury is implicated in the pathogenesis of fibrosis. We identified transitional cells in uninjured *Cldn18* knockout (KO) mouse lungs distinct from previously reported damage-associated transitional progenitors (DATPs) with a less fibrogenic transcriptomic profile. *Cldn18* KO mice are protected from bleomycin-induced fibrosis, with early restoration of cellular homeostasis. Lineage tracing implicates accelerated differentiation as a mechanism for protection from fibrosis, leading us to name these cells regeneration-associated transitional progenitors (RATPs). Multiome confirms that RATPs and DATPs are epigenetically distinct, with RATPs comprised of RATP2s and RATP1s based on epigenomic proximity to AT2s and AT1s, respectively, and suggests dynamic regulatory remodeling during AT2-to-AT1 differentiation, with NKX2.1 and AP-1 active in early transitions and TEAD factors in later stages. These results reveal an unexpected role for *Cldn18* in regulation of AEC plasticity, while identification of RATPs challenges the notion that persistence of transitional alveolar cells is invariably pathologic.

## 1. INTRODUCTION

The lung alveolar epithelium is comprised of two cell types, type 1 (AT1) and type 2 (AT2) cells. AT1 cells are specialized for gas exchange while AT2 cells produce surfactant, are capable of self-renewal and serve as the primary progenitors for AT1 cells at homeostasis and in response to injury. Recently, independent single cell RNA-sequencing (scRNA-seq) studies in murine injury models have identified cells intermediate between AT2 and AT1 cells (Choi et al., 2020; Kobayashi et al., 2020; Strunz et al., 2020), which were variably defined and identified by the expression of combinations of gene markers, namely, pre-alveolar type 1 transitional cell state (PATS; KRT8, KRT19, LGALS3, CLDN4) (Kobayashi et al., 2020), damage-associated transient progenitors (DATPs; KRT8, CLDN4, AREG, KRT7) (Choi et al., 2020), and alveolar differentiation intermediates (ADI; KRT8, CLDN4, TP53) (Strunz et al., 2020). Hereafter, we will refer to them collectively as DATPs.

In mouse models of physiological regeneration such as lipopolysaccharide (LPS)-induced lung injury, AT2 cells exit the cell cycle and adopt a transient transitional state before then differentiating into AT1 cells (Jansing et al., 2017). In contrast, in the bleomycin model of lung injury, alveolar transitional cells marked by KRT8 amongst others, *i.e.* DATPs, accumulate in association with fibrotic regions suggesting that they may be pathologic (Strunz et al., 2020). In human lung, transitional cells have been observed in idiopathic pulmonary fibrosis (IPF) and fibroproliferative acute respiratory distress syndrome (ARDS) (Bharat et al., 2020) and have been termed aberrant basaloid cells, based on their expression of some airway basal cell markers, (*e.g.*, KRT17^+^) (Adams et al., 2020; Habermann et al., 2020; Kathiriya et al., 2022). In both mouse and human, ‘stalled’ transitional cells exhibit cell cycle arrest and are unable to complete the transition to AT1 cells. They express markers of epithelial-mesenchymal plasticity (Adams et al., 2020) and senescence (Auyeung et al., 2022; Strunz et al., 2020), and produce senescence-associated secreted proteins (SASP), key fibrogenic factors implicated in disease pathogenesis.

Of note, in a study on early fibroproliferative human Coronavirus disease 2019 (COVID-19) ARDS, accumulating transitional cells do not express senescence markers and are not associated with fibrosis (Ting et al., 2022), leading the authors to suggest that at least early on, these cells may retain some capacity for physiological differentiation. Although it is possible that the latter transitional cells would have become stalled and fibrogenic if patients had survived longer, this finding also suggests the existence of a potentially non-pathological transient population in early ARDS. More recent studies also suggest alternative transitional states and trajectories between AT2 and AT1 cells, perhaps reflecting stressed vs less stressed states, with more vs less SASP, respectively (Toth et al., 2023; Wang et al., 2023). It therefore appears that different types of transitional cell states may exist, both physiologic and pathologic, and that accumulation of transitional cells may not always be linked to fibrosis.

In mouse, DATPs have been suggested to originate mainly from AT2 cells (Choi et al., 2020; Kobayashi et al., 2020; Strunz et al., 2020), and in both mouse and human (Jaeger et al., 2022; Strunz et al., 2020), a possible airway origin has also been suggested. Several studies have investigated the signaling pathways involved in transitional cell appearance and regulation. WNT/β-catenin signaling has been shown to facilitate lung repair and regeneration following injury (Raslan and Yoon, 2020); however, chronic WNT3A signaling (but not WNT5A or acute WNT3A) induces KRT8 expression and cellular senescence in primary murine AT2 cells *in vitro*, likely promoting AT2-to-DATP transition and impairing transdifferentiation into AT1 cells (Lehmann et al., 2020). Activation of the pro-fibrotic transforming growth factor-β (TGFβ) has also been found to induce the expression of transitional markers in AT2 cells, and inhibition of TGFβ is required for those cells to be able to differentiate into AT1 cells, suggesting that elevated expression of TGFβ in IPF lungs might account for transitional cell persistence (Jiang et al., 2020; Strunz et al., 2020). However, presence of transitional cells during early ARDS without fibrosis (Ting et al., 2022) suggests that a non-TGFβ-dependent/non-fibrogenic transitional cell population may also exist.

Tight junctions (TJ) regulate paracellular permeability and polarity between adjacent epithelial or endothelial cells (Tsukita and Furuse, 2000). In addition to these ‘canonical’ functions, TJs also regulate cell proliferation, differentiation, migration and survival (Spadaro et al., 2012) through mechanisms that include sequestration of mediators and prevention of nuclear translocation of transcription factors (Oka et al., 2010; Zhao et al., 2011). Claudins are integral TJ proteins essential for TJ function (Amasheh et al., 2009; Anderson and Van Itallie, 2009; Rouaud et al., 2020; Suarez-Artiles et al., 2022; Van Itallie and Anderson, 2013). Among twenty-seven claudins (Tamura and Tsukita, 2014), claudin18 (CLDN18) is abundantly expressed in alveolar epithelial cells (AEC), with greater expression in AT1 than AT2 cells (LaFemina et al., 2014; Li et al., 2014). We previously generated mice with global knockout (KO) of both lung-specific (*Cldn18.1*) and stomach-specific (*Cldn18.2*) isoforms (Li et al., 2014). Despite increased barrier permeability, *Cldn18* knockout (KO) mice survive normally, and showed less sensitivity to ventilator-induced lung injury (VILI) (Li et al., 2014). In addition, *Cldn18* KO lungs are enlarged, with expansion and increased proliferation of AT2 cells (Zhou et al., 2018), suggesting a role for TJ in general, and *Cldn18* in particular, in regulation of epithelial progenitor function.

Improved responses of *Cldn18* KO mice to ventilator-induced lung injury (VILI) were attributed to compensatory changes in sodium transport and fluid clearance, but we were curious, given effects of the KO on progenitor function, whether there might also be effects of *Cldn18* loss on AEC differentiation and regeneration following lung injury. Surprisingly, examination of *Cldn18* KO lungs by immunofluorescence revealed the presence of KRT8-and KRT19-expressing alveolar transitional cells at baseline in the absence of fibrosis, suggesting that they differ from previously described DATPs and might not be pathologic. Consistent with this, *Cldn18* KO mice were protected from bleomycin-induced fibrosis, despite the presence of alveolar transitional cells at baseline. ScRNA-seq and multiome analyses identified these as a novel transitional cell population that shows a lower expression of genes involved in stress responses and fibrosis compared to DATPs, and that we named regeneration-associated transitional progenitors (RATPs). *Sftpc*-lineage tracing revealed accelerated differentiation in the KO compared to wild type (WT) mice following injury, suggesting that RATPs are not stalled in their differentiation trajectory. These findings indicate that not all alveolar transitional cells are fibrogenic and suggest that pathways downstream of CLDN18 are key regulators of AEC plasticity.

## 2. METHODS

### 2.1 Animals

Mice were treated in accordance with the Institutional Animal Care and Use Committee (IACUC, USC #21339, UCSD # S21017) guidelines to ensure ethical and humane handling throughout the study. *Cldn18*^−/−^ (KO) mice have been recently reported (Li et al., 2014; Zhou et al., 2018). *Cldn18* KO and WT mice were bred as separate lines and used for experiments as previously described (Li et al., 2014). *Sftpc^+/creERT2^* (Harold A. Chapman, UCSF) mice (Chapman et al., 2011) crossed to *Cldn18^-/-^* mice were further crossed with *ROSA^Tm/Tm^* reporter mice (Arenkiel et al., 2011), yielding mice with the genotype *Sftpc^+/creERT2^;Cldn18^-/-^;ROSA^+/Tm^* for lineage tracing studies. Reporter gene expression was induced by administration of tamoxifen (Tmx) at a dose of 100 mg/kg intraperitoneally (i.p.) for 2 consecutive days. Control mice with the genotype *Sftpc^+/creERT2^; ROSA^+/Tm^* received the same dose of Tmx to label AT2 cells. Tmx clearance was ensured by waiting 30 days before proceeding with bleomycin injury.

### 2.2 Histology and lung fixation

Mouse lungs were perfused with phosphate-buffered saline (PBS), fixed by instillation of 4% paraformaldehyde (PFA) at 20 cmH_2_O pressure, and incubated at 4°C overnight. The following day, PFA was replaced with 70% ethanol and samples were embedded in paraffin. Alternatively, the day following fixation, lobes were placed in PBS followed by a sucrose gradient before cryoembedding in Tissue-tek optimal cutting temperature (Sakura, Torrance, California, USA). Lung sections (5 μm) were prepared using a microtome (Accu-Cut SRM 200, ZEISS, White Plains, New York, USA) and cryostat (CM3050 S, Leica Microsystems, Wetzlar, Germany), respectively, for Sirius red and immunofluorescence staining.

### 2.3 Immunofluorescence staining and image acquisition

Following antigen retrieval in Antigen Unmasking Solution at high pH (Vector Laboratories, Newark, California, USA), lung sections were incubated in CAS block (Invitrogen, Carlsbad, California, USA), and then with primary antibodies overnight at 4° C. Anti-pro-SPC (Seven Hills BioReagents, Cincinnati, Ohio, USA, #9337, 1:500), -AQP5 (Alomone labs, Jerusalem, Israel, #AQP-005, 1:250), -KRT8 Troma 1 (DSHB, Iowa City, Iowa, USA, Supernatant 1mL, 1:30), - KRT19 Troma 3 (DSHB Supernatant 1mL, 1:50), and -tdTomato (SICGEN, Cantanhede, Portugal, #AB8181, 1:50) were used as primary antibodies. Slides were then incubated with rat biotin (Vector Laboratories #BA-9400), Fluorescein avidin (Vector Laboratories, #A-2011), and rabbit 647 (Invitrogen, #A32733), goat 594 (Abcam, Cambridge, United Kingdom, #ab150132), or mouse 647 (Invitrogen, #A32787) Alexa Fluor secondary antibodies. Normal IgGs (Vector), were used as negative controls. Finally, slides were mounted with Vectashield mounting medium including DAPI (Vector). Images were obtained using a BZ-X800 fluorescent microscope (Keyence, Itasca, Illinois, USA) and a Leica SP8 confocal system (Leica Microsystems). Negative controls were used to set the laser intensity for each staining. Quantitation of fluorescence signal was performed using the automated cell quantification BZ-X800 software (Keyence) and QuPath (Bankhead et al., 2017), on five fields from each image.

### 2.4 Sircol assay

Lung tissues were homogenized with a handheld homogenizer in PBS followed by sonication (VirTis, Warminster, Pennsylvania, USA, Virsonic 600) for 45 seconds with a 15 second pause interval 3 times and then centrifuged for 15 min at 2500xg. Supernatants were filtered through 0.45 µm filters (Corning, Corning, New York, USA, #431220). Collagen content was assayed using Sircol soluble Collagen Assay Kit (Biocolor Life Science Assays, Accurate Chemical & Scientific Corporation, Carle Place, New York USA, #S1000).

### 2.5 Bleomycin injury and lung function measurements

Mice were anesthetized with 100 mg/kg of i.p. ketamine (Phoenix Pharmaceuticals, Burlingame, California, USA) and 20 mg/kg of xylazine (Lloyd Laboratories, Walnut, California, USA). Bleomycin (Pfizer, New York, New York, USA, NDC 61703-0332-18) was freshly resuspended and delivered via oral-tracheal administration using a microsprayer (PennCentury, Philadelphia, Pennsylvania, USA). To account for differences in lung size (Zhou et al., 2018), *Cldn18* KO mice were administered 50% greater volume of bleomycin compared to WT, maintaining the same U/body weight (kg) dose. Lung function (compliance and inspiratory capacity) was analyzed using a Flexivent (SCIREQ, Montreal, Ontario, Canada). Following anesthesia, an 18-gauge needle was inserted into the trachea and mice were connected to a Flexivent rodent ventilator. Mice were ventilated under default mechanical ventilation settings (150 breaths/min and tidal volume of 10 ml/kg). Pressure-volume loops were generated to obtain Cst (quasi-static compliance). Cst measurements were performed in each individual subject three times and an average of the three values was further analyzed.

### 2.6 AT2 cell isolation and culture

Primary AT2 cells were isolated from WT and *Cldn18* KO mice as previously described (Demaio et al., 2009). Mouse AEC monolayers (MAECM) were prepared and cultured as previously described (Demaio et al., 2009) using either mouse laminin-1 (Trevigen Minneapolis, Minnesota, USA) or rat laminin-5 (EMD Millipore, Burlington, Massachusetts, USA) to pre-coat Transwell polycarbonate filters (Corning). 10% newborn bovine serum (Thermo Fisher Scientific, Waltham, Massachusetts, USA) was added to Complete Mouse Medium (CMM). Cells were treated at D3 (AT2-like) or D6 (AT1-like) with either 5 mU/ml or 25 mU/ml of bleomycin and harvested for protein isolation and western blotting analysis at D4 and D7, respectively.

### 2.7 Western blot analysis

Proteins were resolved by SDS-PAGE and electrophoretically blotted onto Immune-Blot PVDF membranes. Western blotting was performed using antibodies directed against p-histone H2AX (Cell Signaling, Danvers, Massachusetts, USA, #9718, 1:200) and beta-actin (Abcam, #ab6272, 1:500).

### 2.8 Lung digestion and flow cytometry cell isolation for scRNA-seq

Lungs were harvested at baseline, D3 and D7 following bleomycin from both WT and *Cldn18* KO mice and cell isolation performed as previously reported with minor modifications (Strunz et al., 2020). Briefly, following lung perfusion with PBS, lungs were instilled with and then minced in an enzymatic solution containing dispase (Corning, #354235, 50 caseinolytic units/ml), collagenase (Merck, Rahway, New Jersey, USA, #C1-28, 2 mg/ml), and DNase (QIAGEN, Germantown, Maryland, USA, #79254, 30 µg/ml). Cells were isolated and sorted as CD31^-^/CD45^-^/CD326 (Ep-CAM)^+^ using APC-FIRE CD31 (Biolegend, San Diego, California, USA, #102433, 1:25), BV510 CD45 (Biolegend, # 103137, 1:25) and APC anti-mouse CD326 (Ep-CAM) (Biolegend, #118213, 1:25). To ensure high viability, cells were sorted using the 100 µm nozzle, at a maximal frequency of 1,000 events/sec. Each sample for scRNA-seq consisted of a pool of three biological replicates, of which one mouse carried the lineage tracing for *Sftpc^+/CreERT2^; ROSA^+/Tm^*. To track the source of the isolated cells, each biological replicate was labeled using the anti-mouse hashtag antibody 1-2-3 (Biolegend, #155831, #155833, #155835).

### 2.9 scRNA-seq and data analysis

#### Library preparation, sequencing and quality control

Libraries for scRNA-seq were prepared using the Chromium Next GEM Single Cell 3ʹ Reagent Kit v3.1 (10x Genomics, Pleasanton, California, USA) following the manufacturer’s protocol. Using the software package Cell Ranger v3.0.2 (10x Genomics), sequenced reads were demultiplexed (Cellranger mkfastq) into FASTQ files and aligned (cellranger count) to the mouse reference genome mm10. The expression of *tdTomato* was determined using STARsolo (2.7.9a) (Dobin et al., 2013), aligning raw sequencing reads against a joint *tdTomato* and mm10 reference. *TdTomato* counts, with PCR duplicates removed using unique molecular identifiers (UMIs), were matched to the appropriate cells and incorporated into the Seurat object metadata. Feature barcodes were specified to allow further downstream sample deconvolution based on antibody-derived tags. The filtered count matrices were then loaded into Seurat (Stuart et al., 2019). Likely cell doublets were identified and excluded using two distinct methods. First, cells containing multiple antibody-derived tags were flagged with DropletUtils (Griffiths et al., 2018). Then heterotypic doublets were identified using the scDblFinder v1.13.1 R package (Germain et al., 2021). For initial broad cell type separation, cells with greater than 25% mitochondrial gene content were removed from the overall dataset. Cells were subset into epithelial (*Epcam*^+^) and immune (*Ptprc*^+^) compartments. The *Epcam^+^* population was analyzed for this study. Genes present in at least 3 cells and cells with a minimum of 200 detected genes were included in downstream analysis. Furthermore, cells with greater than 10% mitochondrial gene content were removed from the *Epcam^+^* dataset.

#### Cluster annotation, differential gene expression, and pathway analysis

UMI counts were normalized, scaled, and top variable features identified using the SCTransform (Hafemeister and Satija, 2019) function. UMI matrices were integrated utilizing anchors determined by reciprocal principal component analysis (RPCA). Uniform manifold approximation and projection (UMAP) coordinates and cell clustering were determined by the Seurat functions RunUMAP, FindNeighbors, and FindClusters (principal components 1-30) (Supplemental Figure 1 A). Clusters were annotated using canonical markers (Supplemental Figure 1 B) and downstream differentially expressed gene (DEG) analysis to compare cell populations at different time points and between genotypes was performed using the FindMarkers function in Seurat. Following DEG analysis, Ingenuity Pathway Analysis (IPA, QIAGEN) was performed.

#### RNA velocity

RNA velocity was based on spliced/unspliced gene counts generated by STARsolo (Dobin et al., 2013). RNA velocity was estimated and visualized with velocyto.R (La Manno et al., 2018). The existing UMAP embeddings from the above gene-level analysis were used for visualization of the velocity estimates. Default parameters were used except for the scale value, which was set to “sqrt”.

#### Monocle pseudotime

Pseudotime trajectories were generated using Monocle (v1.3.1) (Cao et al., 2019). Epithelial cells for each genotype were analyzed independently using existing UMAP coordinates, generating principal graphs with default parameters. Root states in the AT2 population were selected manually and used to generate the pseudotime values shown in Figure 7 D.

#### Diffusion pseudotime

To infer trajectory, we performed diffusion pseudotime (DPT) using the Scanpy package (v1.10.3) (Wolf et al., 2018). A neighborhood graph was constructed using sc.pp.neighbors, and the diffusion map was computed with sc.tl.diffmap, using default parameters. CellRank2 (v2.0.7) (Weiler et al., 2024) was used to model and visualize cell transition using the PseudotimeKernel. The resulting pseudotime values and transition matrix were visualized on a diffusion component map.

### 2.10 Multiome

Single-nucleus multiome (RNA + ATAC) libraries were generated and processed with Cell Ranger (10x Genomics) using the mm10 reference genome. WT and *Cldn18* KO datasets were analyzed, and doublets were identified and removed with DoubletFinder. The Seurat object was created from Cell Ranger output and underwent quality control (QC) filtering, retaining nuclei with 1,000 < nCount_ATAC < 100,000, 500 < nCount_RNA < 10,000, 500 < nFeature_RNA < 10,000, mitochondrial transcript percentages (percent.mt) below 5%, and transcription start site enrichment scores (TSSe) greater than 3. After QC filtering and doublet removal, 8,603 high-quality nuclei were retained. To process the multi-modal data, RNA data were normalized using SCTransform, and ATAC data were normalized using TF-IDF and SVD. Multi-modal integration was performed with Seurat functions RunPCA, RunUMAP, FindClusters, and FindMultiModalNeighbors, and peaks were called with MACS2 through Signac. Following initial cell type annotation, epithelial cells were subset, and final clustering was performed with 10 dimensions at resolution 1.5. Marker peaks distinguishing AT2, RATP2, RATP1, and AT1 populations were identified with Seurat’s FindAllMarkers, selecting the top 500 peaks per cell type with average log2 fold change (avg_log2FC) greater than 0.58. ChromVAR was used to identify transcription factor binding motifs, and gene regulatory networks were inferred with SCENIC+, and visualized in Cytoscape.

### 2.11 Statistics

For imaging quantitation using the Automated Cell Quantification BZ-X800 software (Keyence) and QuPath (Bankhead et al., 2017), five random images per sample were quantified and statistical analysis performed. Values are the mean ± SEM. Significance (p < 0.05) for 3 or more group means with 1 and 2 factors was determined by one-way and two-way ANOVA, respectively. Post hoc analyses were performed with Bonferroni’s corrections. Two group means and a 2 × 2 contingency table were compared for significance using 2-sided t tests and Fisher’s exact test, respectively. Z tests were used to determine if ratiometric (i.e., normalized) data were different from control. Statistical analyses for quantitation were performed using GraphPad Prism. For scRNA-seq analyses, Bonferroni’s corrections and Wilcoxon rank sum test were applied using the Seurat package in R Studio. For DEGs, the Fisher’s exact test and Benjamini-Hochberg correction method were applied using IPA.

## 3. RESULTS

### 3.1 KRT8^+^ and KRT19^+^ transitional cells are found in the alveolar region of uninjured *Cldn18* KO mouse lungs in the absence of fibrosis

We previously showed that lungs of *Cldn18* KO mice show increased AT2 cell proliferation (Li et al., 2014). To assess whether *Cldn18* genetic deletion also affects AEC differentiation, we performed immunofluorescence staining for two markers that have been reported to be expressed in alveolar transitional progenitors, KRT8 (Choi et al., 2020; Jiang et al., 2020; Kobayashi et al., 2020; Strunz et al., 2020) and KRT19 (Auyeung et al., 2022). KRT8 and KRT19 were strongly expressed in the airways with minimal expression in the parenchyma of WT mice (Figure 1 A and 1B). In contrast, we detected abundant expression of both KRT8 and KRT19 in the alveolar region of *Cldn18* KO lungs (Figure 1 A and 1 B), indicating transitional AEC expansion at baseline. A significant increase in KRT8 and KRT19 in the alveolar region of *Cldn18* KO lungs was detected at 1 month of age but not at postnatal day 15 and persisted in adult mice (Supplemental Figure 2 A and B). Although several studies in mice as well as in lungs of IPF patients have associated the persistence of transitional AECs variably named ADI/DATP/PATS in the alveolar space with development of fibrosis (Choi et al., 2020; Jiang et al., 2020; Kobayashi et al., 2020; Strunz et al., 2020), surprisingly, Sirius red staining did not reveal fibrosis in *Cldn18* KO lungs at baseline (Figure 1 C and Supplemental Figure 2 C). This was confirmed by similar levels of collagen in *Cldn18* KO and WT lungs by Sircol assay (Figure 1 D). Since fibrosis has been suggested to be due to AECs being stalled in a transitional state (Jiang et al., 2020), we postulated that the absence of fibrosis in *Cldn18* KO lungs might be accompanied by increased ability of transitional AECs to differentiate to AT1 cells.

**Figure 1.**
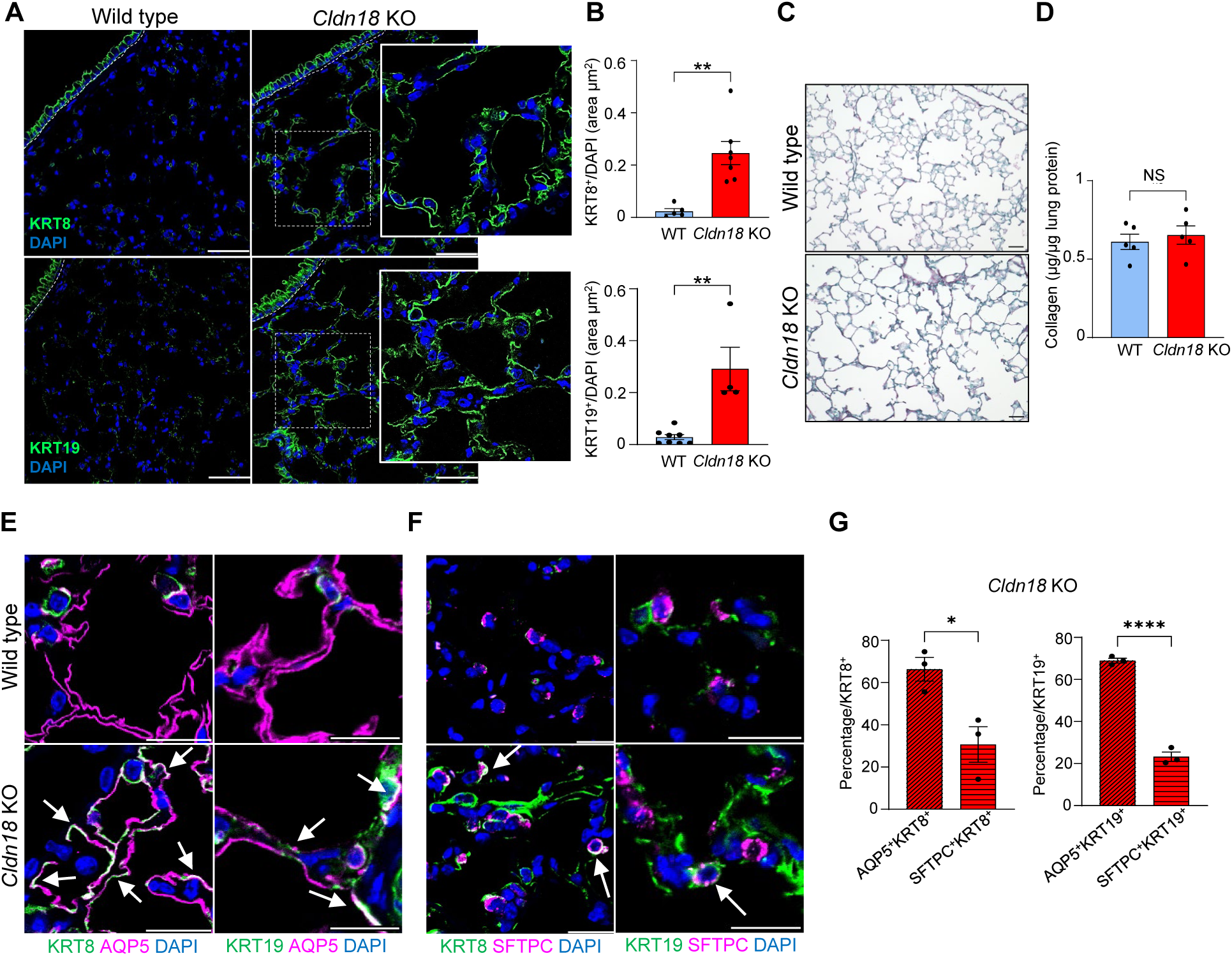
Transitional cells are present in the alveolar region of uninjured *Cldn18* KO mouse lungs without fibrosis. A) Immunofluorescence for KRT8 and KRT19 (green) in *Cldn18* KO and WT lungs of adult (3-6 months old) mice. White dashed lines: airways; white dashed squares: magnified areas shown in the inset; scale bars = 50 µm; n = 6. B) Quantitation of A: top, KRT8^+^, bottom, KRT19^+^ cells over DAPI^+^ area. N = 6, ** = p < 0.01. Unpaired t-test. C) Magnified views from Supplemental Figure 2 C: Sirius red staining of uninjured WT and *Cldn18* KO adult lungs; N = 6, scale bars = 50 µm. D) Quantification of collagen in uninjured WT and *Cldn18* KO adult lungs by Sircol assay; n = 5, NS = not significant. Unpaired t-test. E-F) Magnified views from Supplemental Figure 3 A: E) IF of KRT8 (left) and KRT19 (right) with AQP5 in WT and *Cldn18* KO adult lung. White arrows: co-localization. Scale bars = 20 µm. F) IF of KRT8 (left) and KRT19 (right) with SFTPC in WT and *Cldn18* KO adult lung. White arrows: co-localization. Scale bar = 20 µm. G) Quantitation of E and F; AQP5^+^KRT8^+^ and SPC^+^KRT8^+^ as a percentage of KRT8^+^ (left) and AQP5^+^KRT19^+^ and SPC^+^KRT19^+^ as a percentage of KRT19^+^ (right) in *Cldn18* KO lungs; N= 3; * = p < 0.05; **** = p < 0.001. Unpaired t-test.

To further address whether *Cldn18* KO transitional AECs were on the spectrum between AT2 and AT1 cells, we co-stained the distal lungs of WT and *Cldn18* KO mice for KRT8 and KRT19 together with AT2 and AT1 cell markers. Co-localization of both KRT8 and KRT19 with AQP5 (AT1 cell marker) and SFTPC (AT2 cell marker) was observed in *Cldn18* KO lungs (Figure 1 E and F, and Supplemental Figure 3). In the *Cldn18* KO lungs, ∼70% of KRT8^+^ cells co-expressed AQP5, and ∼30% co-expressed SFTPC; similarly, ∼70% of the KRT19^+^ cells co-expressed AQP5 and ∼23% co-expressed SFTPC (Figure 1 G). On the other hand, among the AQP5^+^ population, ∼15% of cells co-expressed transitional cell markers in the *Cldn18* KO lungs compared to only ∼1% in the WT (Supplemental Figure 3 B and C), while the percentage of WT and *Cldn18* KO SFTPC^+^ cells expressing transitional markers was not different although there was a trend toward an increase in the KO (Supplemental Figure 3 B and C). Thus, the majority of KRT8^+^ and KRT19^+^ cells in the *Cldn18* KO co-expressed AT1 cell markers, indicating transition toward an AT1 cell phenotype while retaining a transitional phenotype. Identification of transitional cells in *Cldn18* KO mice expressing either AT2 or AT1 cell markers suggests a spectrum of differentiation intermediate between AT2 and AT1 cells.

### 3.2 *Cldn18* KO mice are protected from bleomycin-induced lung fibrosis

Given the putative pathogenic role of previously described DATPs in the development of fibrosis, we wondered how the presence of increased transitional cells at baseline in the *Cldn*18 KO might impact the response to fibrotic injury. To address this, we administered bleomycin and examined the lungs of WT and *Cldn18* KO mice at day 21. We initially used a ‘standard’ dose of bleomycin (1.5 U/kg), at which, as expected, WT mice showed significant weight loss (Figure 2 A), mortality (Figure 2 B) and fibrosis (Figure 2 C). Surprisingly, *Cldn18* KO mice showed minimal weight loss, mortality or fibrosis (Figure 2 A-C). We further administered higher doses (2 and 2.5 U/kg) of bleomycin, to ensure that mice were injured. Even at these higher doses, *Cldn18* KO mice were protected from fibrosis as shown by weight, survival and Sirius red staining (Figure 2 D-I). Of note, *Cldn18* KO mice showed 100% survival even at the highest dose of bleomycin (2.5 U/kg), a dose at which 50% of WT mice died by day 14 (Figure 2 H). While lung function was significantly compromised at a dose of 2.5 U/kg of bleomycin in WT mice, neither inspiratory capacity nor compliance were reduced in *Cldn18* KO mice (Figure 2 J, K). To address whether this is due to intrinsic resistance of *Cldn18* KO AT2 cells to bleomycin, we treated AT2 cells isolated from WT and *Cldn18* KO lungs and placed in 2-dimensional culture for 7 days with two bleomycin doses (5 mU/ml and 25 mU/ml), conditions under which AT2 cells transdifferentiate into AT1-like cells by day 6 in culture (Borok et al., 1994). Treatment with bleomycin at day 3 (AT2-like cells) and day 6 (AT1-like cells) led to upregulation of the DNA damage-associated p-histone H2AX 24 hours later (Supplemental Figure 4 A and B), indicating that *Cldn18* KO cells are indeed susceptible to bleomycin. These findings demonstrate that *Cldn18* KO mice are protected from bleomycin-induced fibrosis suggesting that, in contrast to previously reported DATPs, alveolar transitional cells in *Cldn18* KO mice may not be pathologic.

**Figure 2.**
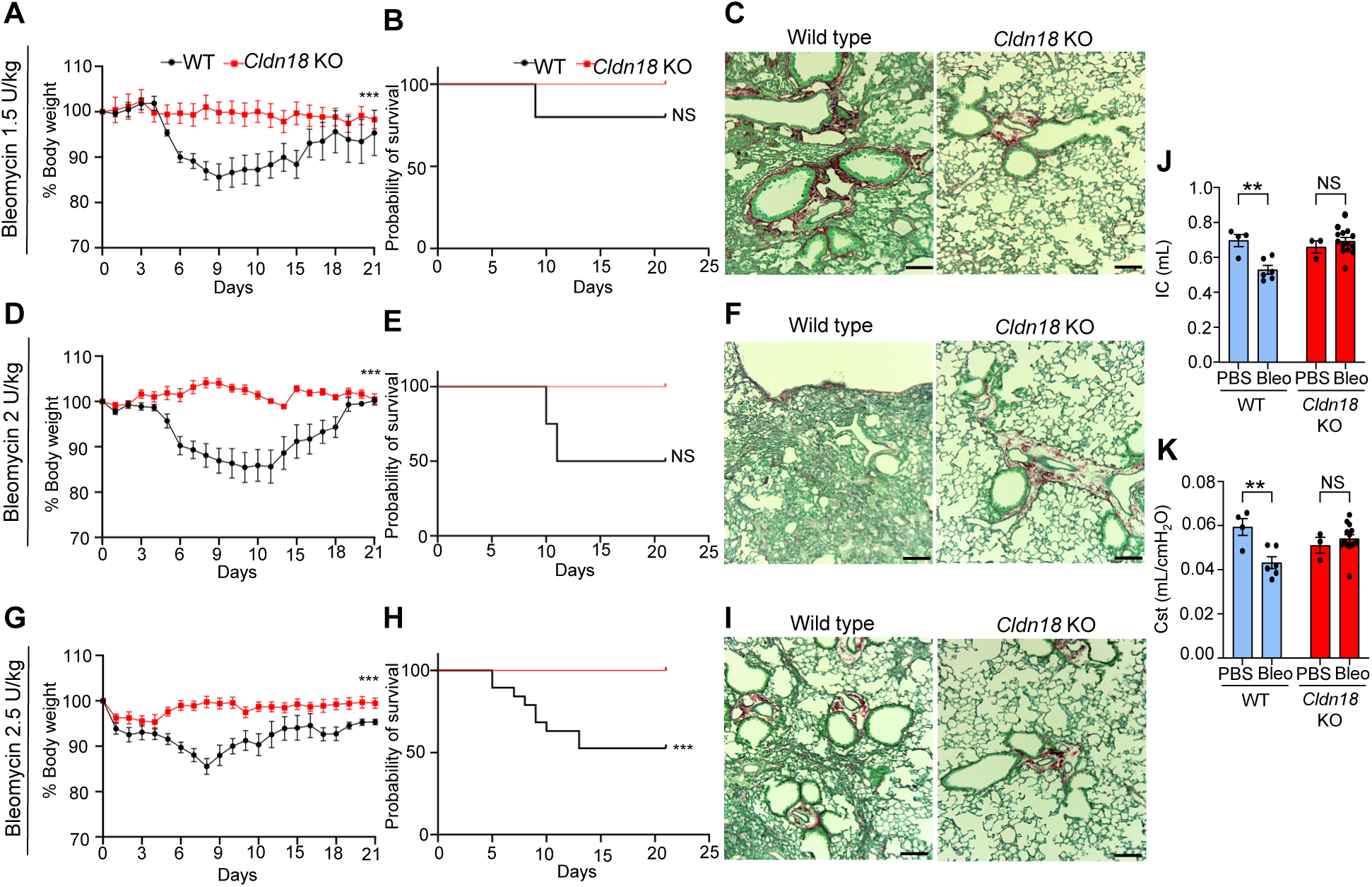
*Cldn18* KO mice are protected from bleomycin-induced pulmonary fibrosis. Bleomycin injury using 1.5 (A-C), 2 (D-F) and 2.5 (G-I) U/kg. Body weight (A, D, G) and survival (B, E, H) over 21 days following bleomycin orotracheal instillation. N = 5 for 1.5 U/kg and 2 U/kg; N=20 for 2.5 U/kg; *** = p<0.005; NS = not significant. Two-way ANOVA. C) Sirius red staining at day 21 following bleomycin. N = 5 for 1.5 U/kg and 2 U/kg; N=20 for 2.5 U/kg, scale bars = 50 µm. D-E) Pulmonary function at day 21 after bleomycin (2.5 U/kg) treatment; J) Inspiratory capacity (IC); K) Static compliance (Cst); n = 4-14; * = p <0.05; ** = p <0.01; *** = p <0.005. Two-way ANOVA.

### 3.3. ScRNA-seq reveals altered cellular composition in *Cldn18* KO lungs at baseline and the presence of alveolar transitional cells

To investigate the composition of the epithelial cells comprising the distal lung and further characterize the alveolar transitional cell population identified in *Cldn18* KO mice, we undertook scRNA-seq of CD45^-^/CD31^-^/EpCAM^+^ cells in *Cldn18* KO and WT mice. Since DATPs have been reported to emerge in WT lungs by day 7-10 following bleomycin injury (Strunz et al., 2020), in addition to baseline, we also performed scRNA-seq on cells isolated from WT and *Cldn18* KO lungs on early days following bleomycin (day 3, D3; day 7, D7) and integrated the data from all days. Integrated UMAP shows that alveolar and airway epithelial cell populations were captured for both WT and *Cldn18* KO samples (Figure 3 A and Supplemental Figure 5). Several previously described clusters, including AT1, AT2 and transitional AECs were identified as defined by markers shown in Supplemental Figure 1 B. Interestingly, striking differences in AEC composition between WT and *Cldn18* KO were observed at baseline (Figure 3 B and C). The major difference noted at day 0 was expansion of the transitional cell population which was increased ∼ three-fold in the *Cldn18* KO compared to WT (Figure 3 B and C), consistent with IF findings (Figure 1). Additional transitional markers were also identified (*i.e., Cldn4, Lgals3, Cd81, Cryab, Clu*) (Supplemental Figure 1 B). Within the AT2 cell population, identified by the expression of known AT2 cell markers (*Sftpc*, *Lamp3*, *Abca3*), we identified five subpopulations: primed AT2 (*Lcn2*, *Lrg1*), AT2-*Lyz1*^+^ (*Lcn2*, *Lrg1*, *Lyz1*), inflamed AT2 (*Lcn2*, *Lrg1*, *Ifitm3*) and proliferating AT2 (*Mki67*) (Supplemental Figure 1 B). We also noted a striking difference in the *Lyz1*^+^ AT2 cell population which represented only 1.12 +/-0.56% of total AECs in the *Cldn18* KO vs 13.36 +/-0.75% in the WT (Figure 3 B-D). There are not extensive studies elucidating the function of *Lyz1*^+^ AT2 cells in lung; however, *Lyz1* has been reported to be associated with AT2 cell maturation (Besnard et al., 2011; Xu et al., 2012). Finally, while most of the AT1 cell markers (*i.e., Rtkn2, Ager, Aqp5, Pdpn, Cav1*) are expressed at similar levels in WT and *Cldn18* KO AT1 cell populations, *Igfbp2*, a marker associated with terminal differentiation (Wang et al., 2018), was virtually absent in *Cldn18* KO AECs (Figure 3 E). These data confirm an increase of alveolar transitional cells as well as differences in cell populations and gene expression that suggest that both AT2 and AT1 cells in the lungs of *Cdn18* KO mice may be less mature.

**Figure 3.**
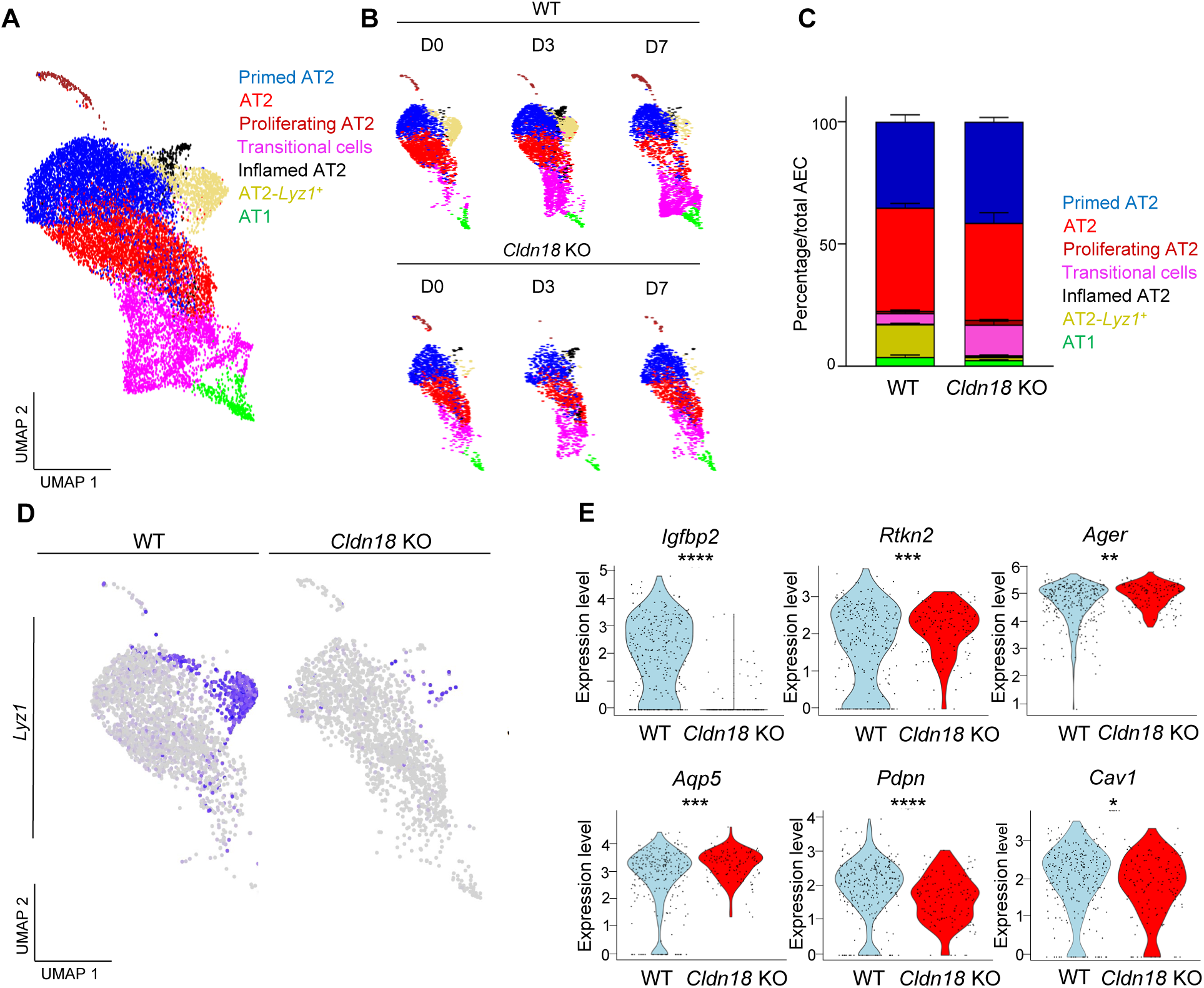
Single cell RNA-seq shows altered cellular composition in *Cldn18* KO lungs at day 0. A) Combined UMAP of WT and *Cldn18* KO alveolar epithelial cells (AEC) at D0. B) UMAPS of WT and *Cldn18* KO AEC populations separated by day. C) Percentages of AEC populations in WT and *Cldn18* KO lungs at baseline. D) UMAP of *Lyz1* expression in WT and *Cldn18* KO AEC. E) Violin plots for gene expression of AT1 cell markers in WT vs *Cldn18* KO AT1 cells. * = p < 0.05, **= p < 0.01, *** = p < 0.001, **** = p < 0.0001. Unpaired t-test.

### 3.4 Single cell RNA-seq identifies a novel population of alveolar transitional cells distinct from DATPs

Given the protection from fibrosis observed in *Cldn18* KO lungs (Figure 2), we hypothesized that the *Cldn18* KO alveolar transitional cells might be fundamentally different from previously described ‘stalled’ and fibrogenic DATPs in WT mice following injury, and instead still capable of differentiation following injury (Choi et al., 2020; Kobayashi et al., 2020; Strunz et al., 2020). Further analysis of the integrated scRNA-seq dataset that includes baseline (day 0) as well as day 3 and day 7 following bleomycin from both WT and *Cldn18* KO, demonstrated that the transitional cell population is comprised of two subsets (Figure 4 A), one resembling previously described DATPs and a second population that we named regeneration-associated transitional progenitors (RATPs) and that is the most prominent at baseline in the lungs of the *Cldn18* KO mice (Figure 5 C-E). The expression of markers commonly used to identify DATPs (Choi et al., 2020; Kobayashi et al., 2020; Strunz et al., 2020) differed between the two alveolar transitional cell subpopulations; for example, *Cldn4*, *Lgals3* and *Sfn* were highly expressed in DATPs, but lower or absent in RATPs (Figure 4 B and Supplemental Figure 6). Differential gene expression analysis showed that among the most significantly upregulated genes in RATPs vs DATPs were markers of AT2 (*i.e., Sftpd, Sftpc, Lamp3*) and AT1 cells (*i.e., Ager, Aqp5*) cells (Figure 4 B and C and Supplemental Figure 6). Furthermore, we identified additional differentially expressed genes between RATPs and DATPs (Figure 4 D), supporting that these are distinct populations. Among these, we found *Trp53*, *Krt17*, and *Pdlim7* to be specifically expressed in DATPs, and *Cited4*, *Bmp4* and *Cox6b2* to be more highly enriched in RATPs (Figure 4 D). Finally, we integrated our dataset with datasets from three other mouse models: LPS (Riemondy et al., 2019), organoids (Kobayashi et al., 2020), and long-term bleomycin (Strunz et al., 2020). While DATPs in our dataset corresponded to those identified in other studies, RATPs localized to a subset of the ‘AT2’ cluster in external datasets (Supplemental Figure 7). Upon integration with our dataset, which encompasses RATPs, this population emerged as a distinct AT2 subcluster, suggesting it may have been previously overlooked and instead annotated as AT2s.

**Figure 4.**
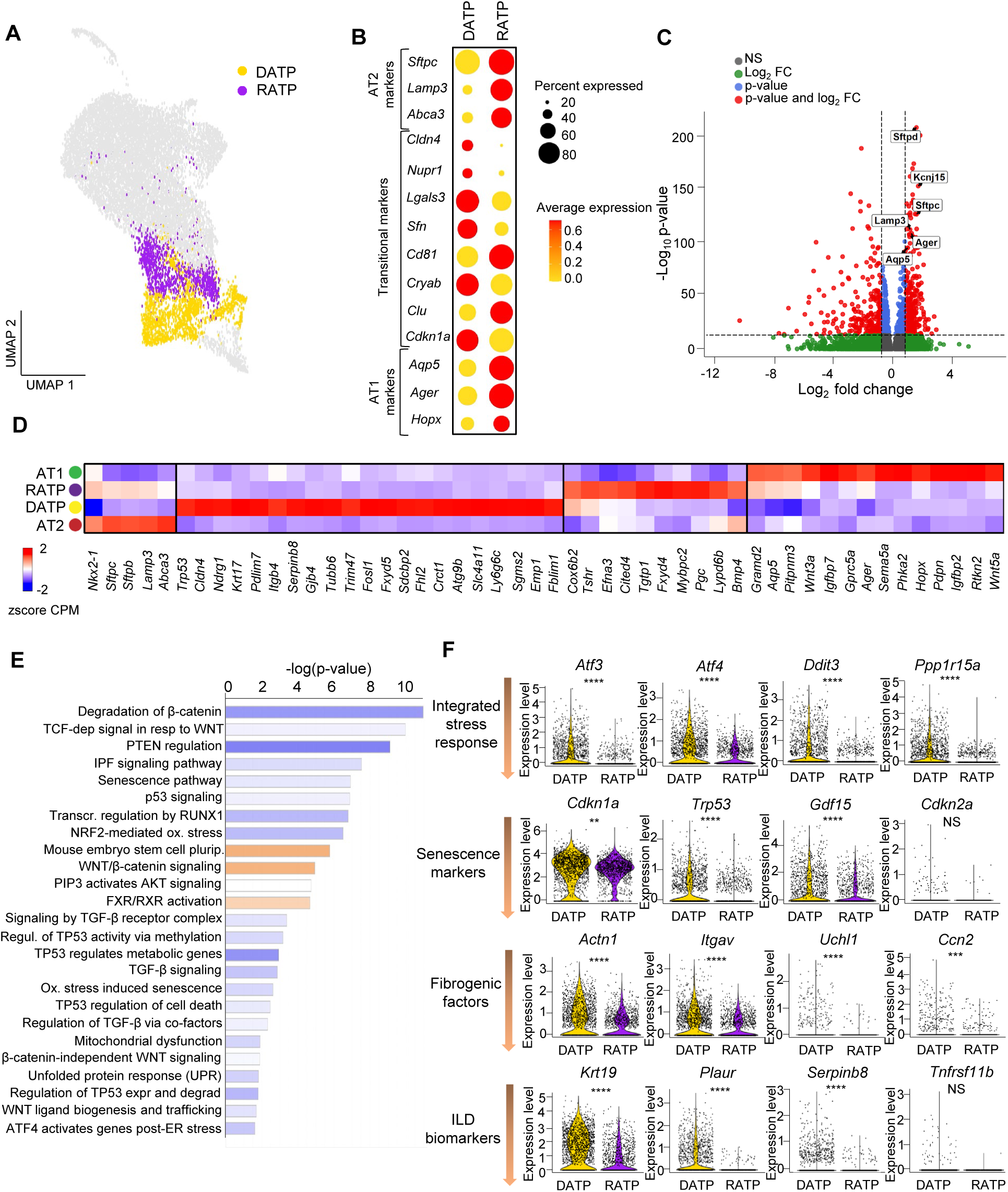
Single cell RNA-seq reveals regeneration-associated transitional progenitors (RATPs) distinct from DATPs. A) Integrated UMAP of all samples at D0, D3 and D7 following bleomycin showing RATPs and DATPs. B) Expression of AT2, DATP and AT1 cell markers in RATPs vs DATPs. C) Volcano plot of gene expression in RATPs vs DATPs. D) Heatmap of gene set enrichment in RATPs, DATPs, AT2 and AT1 cells. E) Ingenuity pathways analysis (IPA) of differential gene expression of RATPs vs DATPs. Blue: inhibited pathways in RATPs; orange: activated pathways in RATPs. F) Violin plots showing markers of integrated stress response, senescence, fibrogenic factors and interstitial lung disease (ILD) biomarkers in RATPs vs DATPs.**= p < 0.01, *** = p < 0.001, **** = p < 0.0001, NS = not significant. Unpaired t-test.

### 3.5 RATPs have a less pathologic transcriptomic profile than DATPs

Several genes used as DATP markers have been implicated in independent studies in IPF pathogenesis, and interestingly we found many of those to be downregulated in RATPs compared to DATPs. *Lgals3*, reported to promote TGFβ1 signaling and fibrosis (Calver et al., 2024; Nishi et al., 2007) and *Sfn*, *Cryab* and *Cdkn1a*, known senescence-associated genes (Arakawa et al., 2022; Lewinska et al., 2017; Limbad et al., 2022; Tamura and Tsukita, 2014), were all elevated in DATPs compared to RATPs (Figure 4 B and Supplemental Figure 6 B), suggesting that RATPs may be less fibrogenic than DATPs. Ingenuity pathway analysis (IPA) comparing differentially expressed genes between RATPs and DATPs indeed revealed that the ‘IPF signaling pathway’, ‘senescence’ and ‘p53’ pathways, as well as ‘TGF-beta’ signaling and ‘unfolded protein response (UPR)’ are all predicted to be reduced in RATPs, while pathways associated with regeneration (‘WNT/beta-catenin’, ‘FXR/RXR activation’) as well as ‘mouse embryonic stem cell pluripotency’ are predicted to be activated (Figure 4 E). One of the most widely adopted markers used to annotate DATPs, the transcription factor nuclear protein 1 (NUPR1), also known to be activated by cellular stress (Huang et al., 2021) and reported to be upregulated in IPF (Yeo et al., 2024), was not only downregulated in RATPs compared to DATPs (Figure 4 B and Supplemental Figure 8), but also predicted to be reduced by IPA upstream analysis, based on the expression of its downstream targets, supporting that RATPs may be less stressed than DATPs. More detailed analysis of the expression profiles of genes implicated in the integrated stress response, ISR (*i.e., Atf3, Atf4, Ddit3, PPp1r15a*) (Huang et al., 2021) and senescence (*i.e., Cdkn1a, Trp53, Gdf15, Cdkn2a*) (Mijit et al., 2020; Radwanska et al., 2022; Saul et al., 2022), fibrogenic factors (*i.e., Actn1, Itgav, Uchl1, Ccn2*) (Anderson and Van Itallie, 2009; Epshtein et al., 2023; Tam et al., 2021; Zhang et al., 1996) and idiopathic lung disease (ILD) biomarkers (*i.e., Krt19, Plaur, Serpinb8, Tnfrsf11b*) (Bowman et al., 2022) found these all to be downregulated in RATPs compared to DATPs (Figure 4 F), reinforcing a less stressed and less fibrogenic phenotype in RATPs compared to DATPs. Together these results support that alveolar transitional cells comprise two subpopulations, of which RATPs have a less pathologic transcriptomic profile than DATPs.

### 3.6 Accelerated restoration of homeostasis and dynamic alterations in cell composition following bleomycin injury in *Cldn18* KO lungs

To better understand the cellular basis for protection in *Cldn18* KO mice and compare dynamic changes in DATPs vs RATPs in WT vs *Cldn18* KO, we analyzed changes in cell composition over time following bleomycin injury. Principal component analysis (PCA) performed on pseudobulk RNA-seq showed that biological replicates cluster closely together (Figure 5 A) demonstrating reproducibility. This was confirmed by the overlap of dimensional plots of biological replicates (Supplemental Figure 9 A) and similar levels of expression of AEC markers (*i.e.*, *Aqp5*, *Sftpc*, *Krt8*) in the biological replicates (Supplemental Figure 9 B). While WT D7 samples clustered separately from WT D0, *Cldn18* KO D7 clusters were close to the *Cldn18* KO D0 (Figure 5 A), suggesting that homeostasis in *Cldn18* KO samples was restored by D7 following injury. Differential gene expression analysis for AEC within the same genotype over time, filtered for significance (adjusted p value ≤0.05), showed that while a similar number of genes was altered in all WT and *Cldn18* KO AECs when comparing D3 vs D0 and D7 vs D3 (Supplemental Figure 10), in the *Cldn18* KO D7 vs D0, only a small number of genes (between 1 and 30) was altered, again supporting return to homeostasis by day 7 following injury (Supplemental Figure 10). Marked differences in the dynamics of cell population composition were observed over time (Figure 5 B, C and D). Transient expansion of inflamed AT2 cells (Figure 5 B and Supplemental Figure 11 A), as well as transient expression of inflammatory markers (*i.e.*, *Isg15*, *Ifitm3*, *Ly6a*) in both WT and *Cldn18* KO AEC populations at D3 post-bleomycin (Supplemental Figure 11 B), demonstrates susceptibility to injury in both genotypes.

**Figure 5.**
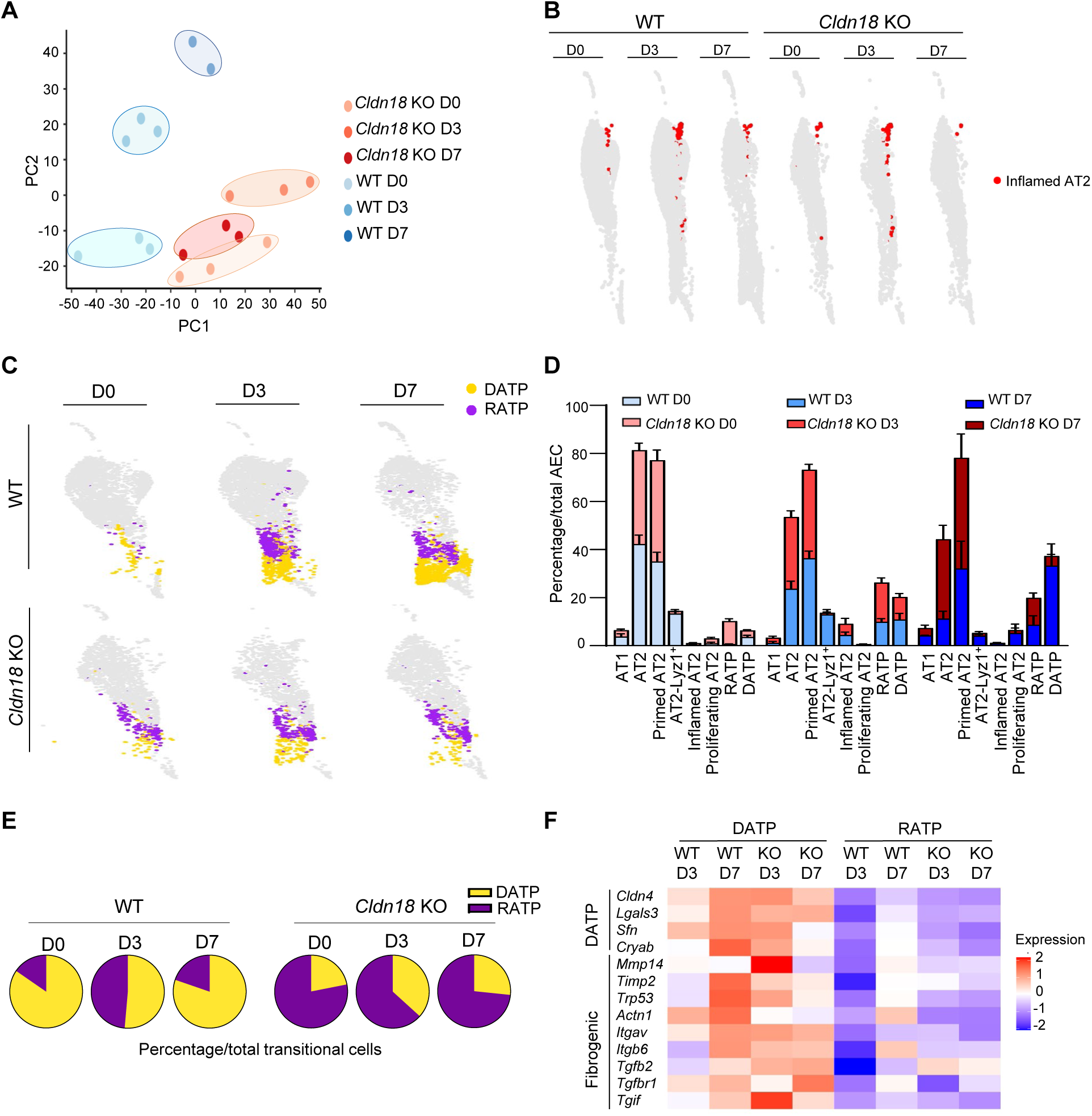
Single cell RNA-seq shows dynamic changes in cell composition following bleomycin injury with earlier restoration of homeostasis in *Cldn18* KO mice. A) Pseudobulk analysis of scRNA-seq. B) UMAP showing inflamed AT2 cells in red. C) UMAP split over time showing DATPs and RATPs in WT and *Cldn18* KO. D) Percentages of AEC subpopulation frequency over time. E) RATPs vs DATPs as a percentage of total transitional AECs. F) Heatmap of DATP and fibrogenic genes in DATP and RATP populations over time following injury.

Changes in transitional cell populations were also observed over time. While in the *Cldn18* KO lungs, abundant RATPs are present at D0 and persist over time, they only expand in the WT at D3 (Figure 5 C). As shown in Figure 5 D, AT2 cells were reduced at D3 in both WT and *Cldn18* KO compared to baseline. By D7, the percentage of AT2 cells reverted to baseline levels in the *Cldn18* KO but was even further reduced in the WT. AT2-*Lyz1*^+^ cells were present in WT lungs at all time-points but did not appear in the *Cldn18* KO lungs, suggesting that the *Cldn18* KO retained a less mature and perhaps more plastic AEC phenotype. Finally, the greatest changes were seen in relative percentages of RATPs and DATPs over time (Figure 5 C, D and E). With respect to total transitional AECs, rare WT transitional cells at D0 are mostly DATPs, with only 15% being RATPs, while in the *Cldn18* KO, RATPs constitute 78%. At D3, RATPs are 49% of total transitional AEC in the WT, compared with 63% in *Cldn18* KO lungs (Figure 5 E). At D7, RATPs decline to 20% in the WT, while they constitute 73% of total transitional AEC in the *Cldn18* KO. DATPs, on the other hand, appeared in the *Cldn18* KO in response to injury at D3 but did not further expand, rather declined by D7, whereas in the WT lungs, they continued to expand over time (Figure 5 C, and D and E). Moreover, expression of fibrogenic genes remained elevated in DATPs relative to RATPs in both WT and *Cldn18* KO lungs at all time-points following injury (Figure 5 F), indicating that, unlike DATPs, RATPs do not adopt a fibrogenic phenotype even in the injured lung.

### 3.7 RATPs are comprised of two subpopulations: RATP1 and RATP2

Since both AT1 and AT2 cell markers are expressed in the RATP population, we wondered whether these markers were co-localized to the same cell or whether RATPs were instead comprised of two separate subpopulations. Staining of *Cldn18* KO lungs at D0 showed that the KRT8^+^AQP5^+^ and KRT8^+^SFTPC^+^ populations were indeed distinct and non-overlapping (Figure 6 A). Reannotation of clusters revealed two subpopulations, with RATP1s expressing mainly AT1 cell markers and RATP2s mainly AT2 cell markers (Figure 6 B and 6 C and Supplemental Figure 12). To better define these subpopulations, we evaluated the relative expression of AT2, AT1 and DATP markers in all the transitional cell populations (Figure 6 C). When compared to either RATP1s or DATPs, the RATP2 population was characterized by expression of the AT2 cell markers (*i.e.*, *Sftpd, Sftpb*) (Figure 6 C). In contrast, AT1 cell markers (*i.e, Ager, Aqp5, Hopx, Rkn2, Gprc5a, Gramd2*) were more highly expressed in RATP1s compared to either RATP2s or DATPs. To determine whether both populations differ from DATPs, we analyzed the expression of genes involved in fibrosis, SASP, DNA damage, cell death, and proteostasis (Wang et al., 2023) separately in RATP1s and RATP2s, and found that these were all expressed at lower levels in RATP1s and RATP2s, indicating that both populations are less fibrogenic and less stressed than DATPs (Figure 6 D).

**Figure 6.**
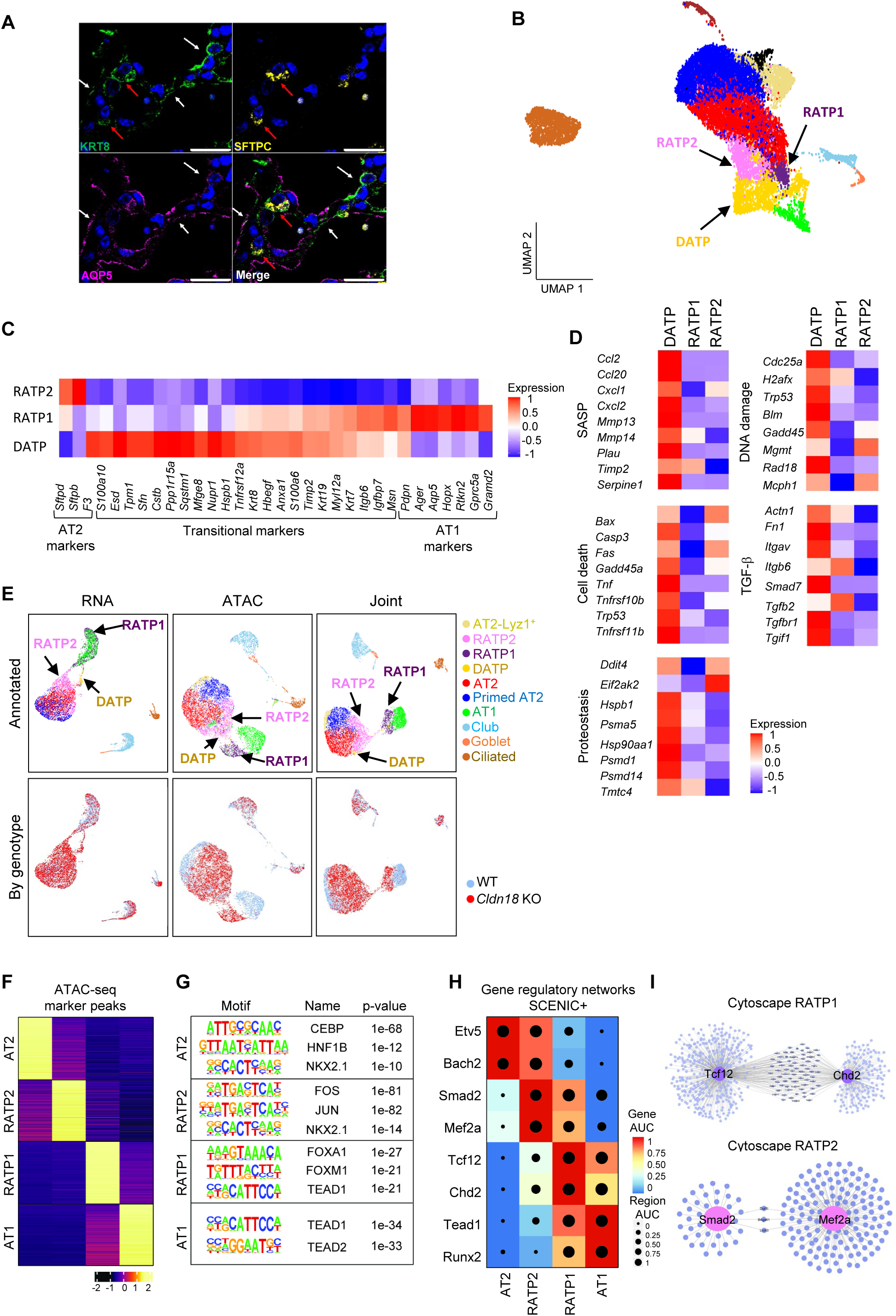
RATPs are comprised of two subpopulations, RATP1s and RATP2s. A) Triple staining for AQP5, SFTPC and KRT8; N= 4; scale bars = 20 µm. White arrows: co-localization of KRT8 and AQP5; red arrows: co-localization of KRT8 and SFTPC. B) UMAP with RATP1, RATP2, and DATP annotation. C) Expression of AT2, transitional and AT1 markers in DATP, RATP1 and RATP2 cell populations. D) Expression of SASP, cell death, DNA damage, proteostasis and TGF-b markers in DATPs, RATP1s and RATP2s. E) snMultiome: Top: UMAPs show cell clustering by RNA and ATAC features separately, and jointly (WNN). Cell type annotations matching to single cell annotations. Bottom: Clustering by genotype. F) Heatmap of the top 500 marker ATAC-seq peaks for each cell state (AT2, RATP2, RATP1, AT1) showing normalized accessibility (z-score). Peaks were identified using differential accessibility analysis across the four populations. G) Top enriched transcription factor motifs identified within cell state-specific accessible peaks for each population. Representative motifs and their associated transcription factors are shown with enrichment p-values. H) Gene regulatory network activity inferred using SCENIC+ across populations. Heatmap shows gene AUC (color) and region AUC (dot size) for selected regulons active across AT2, RATP2, RATP1, and AT1 populations. I) SCENIC+ gene regulatory network highlighting connections among candidate regulators and their predicted target genes in RATP populations. Node size reflects the number of predicted targets and edges indicate inferred regulatory connections.

### 3.8 Multiome further resolves RATP states and identifies cell-specific gene regulatory networks

To investigate the gene regulatory mechanisms underlying the accumulation of RATP2 and RATP1 populations in *Cldn18* KO lungs, we performed single nucleus multiome (RNA + ATAC) profiling of *Cldn18* KO and WT lungs at baseline. Integration of the transcriptome and chromatin accessibility showed that RATP1 and RATP2 segregate out readily, confirming scRNA-seq findings and extending beyond that, with very few DATPs at baseline (Figure 6 E). The joint UMAP shows that RATP2s cluster close to AT2s, while RATP1s cluster close to AT1s, suggesting a spectrum from AT2>RATP2>RATP1>AT1. Furthermore, the joint UMAP further confirmed that RATPs were mainly present in the KO (Figure 6 E). We next defined cell state-specific chromatin accessibility profiles, identifying marker ATAC peaks distinguishing AT2, RATP2, RATP1, and AT1 populations (Figure 6 F). Marker peaks were highly cell-specific, supporting that AT2, RATP2, RATP1, and AT1 are epigenetically distinct cell states. Transcription factor motif enrichment analysis of cell state marker peaks revealed enrichment of CEBP, HNF1B, and NKX2.1 motifs in AT2 cells, and AP-1 family motifs (FOS, JUN) in RATP2s, and FOXA1, FOXM1, and TEAD1 motifs in RATP1s, with TEAD motifs persisting into AT1 cells (Figure 6 G). These patterns suggest dynamic remodeling of the regulatory landscape during AT2-to-AT1 cell differentiation, with NKX2.1 and AP-1 activity marking early transitional states and TEAD factors associated with later transitions.

To better understand transcriptional control underlying the RATPs, we applied SCENIC+, a tool that infers the activity of transcription factors and their target genes (‘regulons’) from scRNA-seq data. This analysis revealed that distinct cell populations (AT2, RATP2, RATP1, AT1) are each defined by specific sets of active gene regulatory programs (Figure 6 H). In particular, RATP2s and RATP1s showed increased activity of regulons controlled by *Smad2*, *Mef2a*, *Tcf12*, *Chd2*, and *Tead1*, suggesting that these transcription factors play key roles in sustaining the transitional states that expand in the *Cldn18* KO lungs.

Finally, we reconstructed regulatory networks highlighting Smad2 and Mef2a for RATP2s and Tcf12 and Chd2 for RATP1s as candidate transcriptional regulators with broad target gene connectivity in RATP populations, as shown by Cytoscape analyses (Figure 6 I), a platform for visualizing and analyzing molecular interaction networks. These findings suggest a coordinated network driving the accumulation of regenerative intermediates upon loss of *Cldn18*.

### 3.9 Accelerated differentiation in *Cldn18* KO lungs following injury

The presence of non-fibrogenic transitional AECs expressing AT1 cell markers in *Cldn18* KO mice suggested that RATPs are not stalled and retain the capacity to transition into AT1 cells. To further investigate this, we performed trajectory analysis on our integrated scRNA-seq dataset. We organized cells using diffusion maps (Figure 7 A) and performed pseudotime analysis (Figure 7 B) for both WT and *Cldn18* KO. Interestingly, while in the WT, the trajectory branched into two terminal points, AT1s and DATPs, in the *Cldn18* KO, the trajectory culminated only in the AT1 population (Figure 7 A and B). Distribution of individual cells according to their cell identities and their calculated pseudotime suggested that there are two terminal stages in the WT (DATP and AT1), with only one represented by AT1s in the *Cldn18* KO (Supplemental Figure 13 A). Single-cell trajectory analysis using Monocle (Trapnell et al., 2014), which predicts the cell-to-cell relationship in pseudotime and is able to ‘learn’ if cells should be placed in the same trajectory or different trajectories, showed similar trajectories to those described above (Figure 7 C and D). Interestingly, Monocle trajectory in the *Cldn18* KO passes through the RATP2 and RATP1 populations but not the DATPs, supporting that RATPs are the main source of AT1 cells in the *Cldn18* KO following injury dataset. Finally, RNAvelocity analysis, which estimates the future state of cells by analyzing the ratio of spliced and unspliced RNA transcripts, suggested that *Cldn18* KO RATPs are predicted to transition to AT1 cells (Supplemental Figure 13 B). Thus, independent trajectory analyses suggest that the RATP population present in the *Cldn18* KO lungs is capable of transdifferentiating into AT1 cells already by D7 following bleomycin injury and that this trajectory predominates over the ‘stalled’ DATP population.

**Figure 7.**
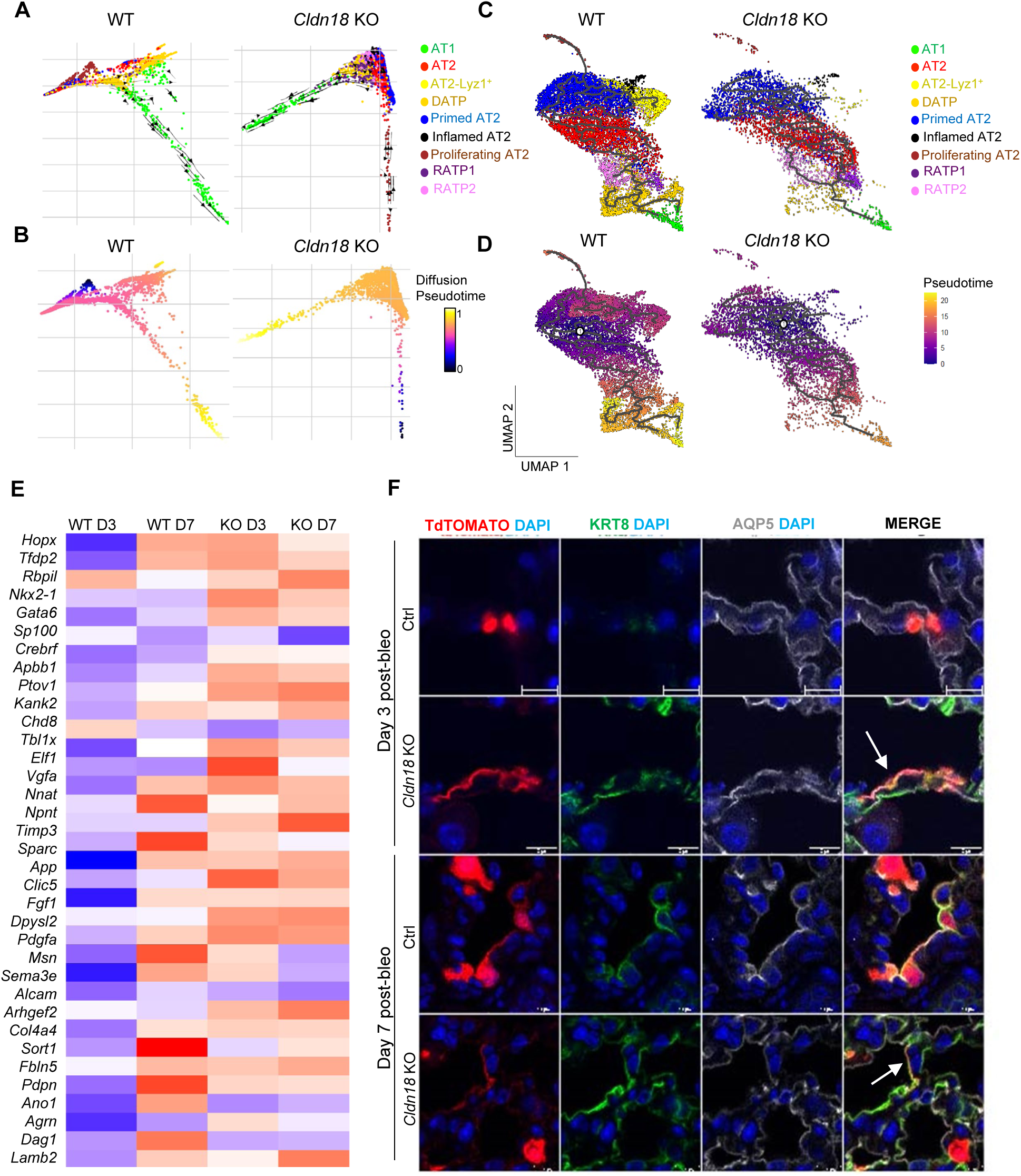
Differentiation is accelerated in *Cldn18* KO compared to WT lung following bleomycin injury. A) Diffusion map showing cell annotations. B) Diffusion map showing pseudotime. C) Monocle analysis showing cell populations. D) Monocle analysis showing pseudotime. E) Heatmap showing expression of ‘AT1 terminal’ genes as defined by Strunz *et al*., in RATPs. F) Triple staining for tdTOMATO (SFTPC-lineage traced), KRT8, and AQP5 at D3 and D7 following bleomycin injury. N=6, scale bars = 20µm.

To gain further insight into the rate of RATP differentiation toward the AT1 cell fate in *Cldn18* KO compared to WT lungs and investigate whether protection from fibrosis might be due to RATPs being poised for transdifferentiation into AT1 cells, we analyzed the expression of a gene set previously identified by Strunz *et al*. and found to be upregulated in the KRT8^+^ ADI ‘terminal AT1 state’ of differentiation (Strunz et al., 2020). We observed that *Cldn18* KO RATPs exhibited elevated expression of these ‘terminal AT1 state’ genes as early as D3 post-injury, most of which persisted through D7, supporting our hypothesis that alveolar regeneration initiates at very early stages following bleomycin insult in this mouse model (Figure 7 E). In contrast, in the WT, where RATPs expand at D3 post-bleomycin and then become outnumbered by DATPs, expression of the ‘terminal AT1 state’ gene signature was only observed in the remaining RATPs at D7, but not at D3 (Figure 7 E). This temporal difference suggests that RATPs in the *Cldn18* KO may be primed to respond rapidly to injury which ultimately leads to protection from fibrosis.

Finally, to confirm accelerated AT1 cell differentiation in *Cldn18* KO mice, we used lineage tracing, using *Sftpc^+/creERT2^Cldn18^-/-^ROSA^Tm/Tm^* and *Sftpc^+/creERT2^Cldn18^+/+^ROSA^Tm/Tm^* mice and analyzed injured lungs at early time points following bleomycin injury (D3 and D7). The reporter was activated by tamoxifen (TMX) injection (100 mg/kg on two consecutive days) followed by a 30-day washout period before instilling bleomycin (2 U/kg). At D3 following bleomycin, lineage-labeled cells in the *Cldn18* KO lungs assumed an AT1 cell-like shape, suggesting that they are already undergoing transdifferentiation (Figure 7 F). Triple staining of tdTomato with KRT8 and AQP5 showed that lineage-labeled cells at D3 in the *Cldn18* KO lungs express markers of both transitional and AT1 cells (Figure 7 F), supporting the scRNA-seq analysis, and suggesting that lungs of *Cldn18* KO mice exhibit accelerated regeneration following injury. This was further supported by scRNA-seq analysis showing *Tomato* expression in *Cldn18* KO AT1 cells already at D3 following injury (Supplemental Figure 13 C). These data suggest that protection from fibrosis in the *Cldn18* KO lungs is conferred by accelerated regeneration and importantly, that the presence of alveolar transitional cells is not always associated with fibrosis following bleomycin injury.

## DISCUSSION

Recent studies have increasingly highlighted the significant role of pathologic alveolar transitional progenitors in the pathogenesis of lung fibrosis (Choi et al., 2020; Jiang et al., 2020; Kobayashi et al., 2020; Strunz et al., 2020). These cells, characterized by their unique transitional state between AT2 and AT1 cells, express profibrotic and proinflammatory mediators, contributing to the fibrotic niche observed in conditions such as idiopathic pulmonary fibrosis (IPF) (Jiang et al., 2020; Strunz et al., 2020). Consequently, the prevailing notion in the field is that it is the persistence of DATPs in the lungs that inevitably leads to fibrosis. Here, we identified a novel population of transitional AECs that we named regeneration-associated transitional progenitors (RATPs), that is expanded in the lungs of *Cldn18* KO mice at baseline in the absence of fibrosis and is associated with protection from bleomycin injury. RATPs express some previously reported transitional AEC markers (e.g., KRT8, KRT19) but have a less fibrogenic profile despite their persistence at baseline and over time after bleomycin injury. Co-localization of KRT8 and KRT19 with AT1 cell markers in a subset of these cells suggests that *Cldn18* KO RATPs are not ‘stalled’ but rather in a state of transdifferentiation. Consistent with this, scRNA-seq trajectory analyses suggested that *Cldn18* KO RATPs, unlike DATPs, predominantly differentiate into AT1 cells following bleomycin injury.

*Cldn18* KO mice are protected from bleomycin-induced fibrosis, and lineage tracing with *Sftpc*, that could potentially label either AT2 cells or RATP2s, revealed early appearance of the lineage label in *Cldn18* KO cells expressing KRT8 and AQP5 already by D3 following injury. This suggests that accelerated AT2 (or RATP2) to AT1 cell differentiation leads to enhanced regeneration in *Cldn18* KO lungs and that *Cldn18* KO lungs are primed to respond rapidly to injury. Interestingly, scRNA-seq analysis over time after injury showed that RATPs increase in both WT and *Cldn18* KO at D3 following bleomycin, with a greater increase in the KO as a percentage of the total transitional cell population. DATPs on the other hand were similar on D3 in the WT and the *Cldn18* KO but persisted in the WT at D7 and had already returned to baseline in the *Cldn18* KO. Overall, at D7, the relative fraction of RATPs in the total transitional AEC population was high in the *Cldn18* KO and minimal in the WTs while DATPs constituted the biggest portion of transitional AECs in the WT and were far less abundant in the *Cldn18* KO. Although recent studies have suggested alternative transitional states and trajectories between AT2 and AT1 cells, perhaps reflecting stressed vs less stressed states (Toth et al., 2023; Wang et al., 2023), this RATP population has not been identified to date. Our integration with other datasets suggested that this population might have been overlooked and annotated as part of the AT2 population in other studies (Supplemental Figure 7). At a molecular level, further analysis revealed that, in the *Cldn18* KO lungs, the RATP population exhibited high expression of genes identified as markers of the “AT1 terminal” state of transitional cells, as defined by *Strunz et al*., at both D3 and D7 following bleomycin treatment suggesting that in the *Cldn18* KO lungs this population responds in a timely manner to bleomycin injury. Indeed, our study suggests that genetic deletion of *Cldn18* creates a favorable environment for accelerated regeneration as evidenced by lineage tracing and scRNA-seq analyses, ultimately protecting against fibrosis. Lineage tracing of RATPs will be needed to establish the rate of differentiation of these cells at baseline and following injury.

One caveat of our study is that, given the rapid regeneration and absence of fibrosis following bleomycin injury in the *Cldn18* KO lungs, the extent of injury can be difficult to assess. However, several lines of evidence indicate that *Cldn18* KO mice were indeed injured following bleomycin: inflamed AT2 appeared in the *Cldn18* KO lungs as in the WT and inflammatory genes were transiently upregulated in the AEC population at D3 following injury; isolated *Cldn18* KO AT2 cells were susceptible to bleomycin *in vitro*; cell population proportions changed in the *Cldn18* KO lungs following bleomycin injury; and lineage tracing showed rapid responses of labeled cells following bleomycin injury in *Cldn18* KO lungs.

RATPs and DATPs were identified in our scRNA-seq data as distinct populations with some overlap in classical transitional AEC markers (*e.g.*, KRT8 and KRT19). Importantly, RATPs were marked by reduced expression of integrated stress response markers, senescence markers, fibrogenic factors, and interstitial lung disease (ILD) markers compared to DATPs, suggesting that they are not ‘pathologic’ as reported for DATPs. Analysis via Ingenuity Pathway Analysis (IPA) of RATPs vs DATPs revealed activation of WNT/β-catenin signaling, known to promote differentiation of epithelial cells contributing to the repair process (Raslan and Yoon, 2020), in RATPs, and inhibition of TGF-β and TP53 pathways that characterize instead the DATP population (Jiang et al., 2020). DATPs and RATPs remained distinct over time after injury; notably, both RATP1s and RATP2s maintained a non-fibrogenic phenotype compared to DATPs as evidenced by scRNA-seq and appear to be poised for differentiation rather than becoming “stalled” post-injury, as suggested by their expression of “AT1 terminal” genes and pseudotime. Interestingly, *Pdlim7*, a marker that has been associated with a more senescent subset of DATPs (Wang et al., 2023), was expressed in DATPs but not in RATPs in our study. Among markers utilized to identify DATPs (Choi et al., 2020), the genes exhibiting reduced expression in RATPs compared to DATPs were those associated with fibrosis in previous studies. Among these, *Stratifin*, also known as 14-3-3σ (SFN), has been identified as a diagnostic biomarker in serum of drug-induced interstitial lung disease and diffuse alveolar damage (Arakawa et al., 2022) and is overexpressed in IPF lungs (Mayr et al., 2021), while *Nupr1*, a marker and regulator of oxidative stress (Huang et al., 2021), has been recently identified as marker of advanced IPF (Yeo et al., 2024), Thus, RATPs, appear to be less stressed and less fibrogenic than DATPs and associated with accelerated regeneration following injury.

Intriguingly, immunofluorescence co-localization further revealed that the RATP population is comprised of two subpopulations, one expressing AT1 cell markers, that we called RATP1s, and one expressing AT2 cell markers, that we called RATP2s. ScRNA-seq and multiome analyses clearly showed that RATP2s and RATP1s cluster separately, with RATP2s clustering closer to AT2s and RATP1 closer to AT1, suggesting a differentiation trajectory from AT2 to RATP2, RATP1 and finally AT1 that will need to be confirmed by lineage tracing. Importantly, both RATP1s and RATP2s showed downregulation of markers of SASP, cell death, DNA damage, TGFβ, and proteostasis when compared to DATPs.

CLDN18, together with other tight junction (TJ) proteins, regulates paracellular permeability and cell polarity in the lung epithelium (Li et al., 2014), and has been identified as part of a signaling hub that regulates AT2 cell proliferation by modulating the Hippo signaling pathway (Zhou et al., 2018). Although *Cldn18* KO mice survive normally (Li et al., 2014), they exhibit increased solute permeability and alveolar fluid clearance (AFC) compared to WT mice, which conceivably might lead to subclinical injury in the lungs of *Cldn18* KO mice. We speculate that it is this low-grade injury due to TJ disruption that may be responsible for the RATP expansion observed at baseline in the *Cldn18* KO lungs. *Cldn18* KO AECs likely perceive “injury signals” from their surrounding environment that induces them to respond by increasing AT2-to-AT1 transition. This constant turnover may explain, at least in part, the absence of “mature” markers of AECs in the lungs of *Cldn18* KO mice, as evidenced by the lack of *Igfbp2* (Wang et al., 2018) expression in *Cldn18* KO AT1 cells, as well as absence of the *Lyz1*^+^ AT2 cell population. Although the role of *Lyz1* has not been extensively investigated, its expression in the AT2 cell population has been associated with maturation (Besnard et al., 2011; Xu et al., 2012). We speculate that incomplete maturation of the alveolar cells in the *Cldn18* KO lungs contributes to their retention in a more plastic state, which enhances their capacity for adaptive responses and regeneration. Indeed, in various biological contexts, progenitor and stem cells often remain in a partially differentiated state, allowing them to respond effectively to physiological demands and environmental cues (Wabik and Jones, 2015). Furthermore, the expansion of RATPs in the *Cldn18* KO lungs suggests a novel property whereby weakening of TJ proteins, signaling a subclinical baseline injury, could unleash AEC plasticity to drive repair.

In conclusion, our study demonstrates that the presence and persistence of transitional AECs *per se* does not induce fibrosis. Instead, a more regenerative cell population that we called RATPs, emerges in *Cldn18* KO mice and may in fact be primed to respond to injury, potentially linked to impaired maturation due to disruption of epithelial tight junctions, thereby protecting from fibrosis due to enhanced tissue repair. Future studies are needed to interrogate the pathways and signals that allow for RATP expansion in the *Cldn18* KO lungs. We speculate that the subclinical injury imparted by tight junction disruption, together with the increased proliferation of AT2 cells due to activation of the Hippo/YAP signaling that we previously reported in the *Cldn18* KO lungs (Zhou et al., 2018), induces a constant turnover of the AEC population. Consistent with this, SCENIC analysis showed increase of the TEAD gene regulatory network in RATP1s which was further augmented in AT1 cells. Interestingly, the GRN analysis defined *Smad2* and *Mef2a* as regulators of RATP2 populations, consistent with their role in initiating early transitional programs (Bhaskaran et al., 2007; Zhao et al., 2013; Zhou et al., 2021). On the other hand, *Tcf12* and *Chd2* emerge as novel candidate regulators that may contribute to reinforcing chromatin remodeling programs during the late stages of AT1 differentiation, opening the avenue for future studies to experimentally test their specific roles in alveolar maturation and AT1 differentiation. Uncovering the molecular and cellular mechanisms that lead to RATP expansion and accelerated differentiation in *Cldn18* KO mice has the potential to be paradigm-shifting and will be crucial to harnessing AEC plasticity to enhance regeneration following injury, potentially offering new therapeutic strategies for fibrotic lung disease.

## Acknowledgements and funding

This study was funded by the National Heart, Lung, and Blood Institute (NHLBI) R35 HL135747 (ZB), R01HL114959 (BZ) and the NIH/NCI Norris Comprehensive Cancer Center core grant P30CA014089. Histology and microscopy services were provided by the Cell and Tissue Imaging Core of the USC Research Center for Liver Diseases (P30 DK048522) and the UCSD School of Medicine microscopy core (NS047101). The authors thank Ray Juan Alvarez for technical support with AT2 isolation, and Drs. Lucy Mary Golden, Ana Claudia Maretti Garcia and Omar Lakhdari for support with flow cytometry and single-cell library preparation for RNA sequencing.

## REFERENCES

1. Adams, T.S., Schupp, J.C., Poli, S., Ayaub, E.A., Neumark, N., Ahangari, F., Chu, S.G., Raby, B.A., DeIuliis, G., Januszyk, M., et al. (2020). Single-cell RNA-seq reveals ectopic and aberrant lung-resident cell populations in idiopathic pulmonary fibrosis. Sci Adv 6, eaba1983.

2. Amasheh, S., Milatz, S., Krug, S.M., Markov, A.G., Gunzel, D., Amasheh, M., and Fromm, M. (2009). Tight junction proteins as channel formers and barrier builders. Ann N Y Acad Sci 1165, 211–219.

3. Anderson, J.M., and Van Itallie, C.M. (2009). Physiology and function of the tight junction. Cold Spring Harb Perspect Biol 1, a002584.

4. Arakawa, N., Ushiki, A., Abe, M., Matsuyama, S., Saito, Y., Kashiwada, T., Horimasu, Y., Gemma, A., Tatsumi, K., Hattori, N., et al. (2022). Stratifin as a novel diagnostic biomarker in serum for diffuse alveolar damage. Nat Commun 13, 5854.

5. Arenkiel, B.R., Hasegawa, H., Yi, J.J., Larsen, R.S., Wallace, M.L., Philpot, B.D., Wang, F., and Ehlers, M.D. (2011). Activity-induced remodeling of olfactory bulb microcircuits revealed by monosynaptic tracing. PLoS One 6, e29423.

6. Auyeung, V.C., Downey, M.S., Thamsen, M., Wenger, T.A., Backes, B.J., Sheppard, D., and Papa, F.R. (2022). IRE1alpha drives lung epithelial progenitor dysfunction to establish a niche for pulmonary fibrosis. Am J Physiol Lung Cell Mol Physiol 322, L564–L580.

7. Bankhead, P., Loughrey, M.B., Fernandez, J.A., Dombrowski, Y., McArt, D.G., Dunne, P.D., McQuaid, S., Gray, R.T., Murray, L.J., Coleman, H.G., et al. (2017). QuPath: Open source software for digital pathology image analysis. Sci Rep 7, 16878.

8. Besnard, V., Wert, S.E., Ikegami, M., Xu, Y., Heffner, C., Murray, S.A., Donahue, L.R., and Whitsett, J.A. (2011). Maternal synchronization of gestational length and lung maturation. PLoS One 6, e26682.

9. Bharat, A., Querrey, M., Markov, N.S., Kim, S., Kurihara, C., Garza-Castillon, R., Manerikar, A., Shilatifard, A., Tomic, R., Politanska, Y., et al. (2020). Lung transplantation for patients with severe COVID-19. Sci Transl Med 12.

10. Bhaskaran, M., Kolliputi, N., Wang, Y., Gou, D., Chintagari, N.R., and Liu, L. (2007). Trans-differentiation of alveolar epithelial type II cells to type I cells involves autocrine signaling by transforming growth factor beta 1 through the Smad pathway. J Biol Chem 282, 3968–3976.

11. Borok, Z., Danto, S.I., Zabski, S.M., and Crandall, E.D. (1994). Defined medium for primary culture de novo of adult rat alveolar epithelial cells. In Vitro Cell Dev Biol Anim 30A, 99–104.

12. Bowman, W.S., Newton, C.A., Linderholm, A.L., Neely, M.L., Pugashetti, J.V., Kaul, B., Vo, V., Echt, G.A., Leon, W., Shah, R.J., et al. (2022). Proteomic biomarkers of progressive fibrosing interstitial lung disease: a multicentre cohort analysis. Lancet Respir Med 10, 593–602.

13. Calver, J.F., Parmar, N.R., Harris, G., Lithgo, R.M., Stylianou, P., Zetterberg, F.R., Gooptu, B., Mackinnon, A.C., Carr, S.B., Borthwick, L.A., et al. (2024). Defining the mechanism of galectin-3-mediated TGF-beta1 activation and its role in lung fibrosis. J Biol Chem 300, 107300.

14. Cao, J., Spielmann, M., Qiu, X., Huang, X., Ibrahim, D.M., Hill, A.J., Zhang, F., Mundlos, S., Christiansen, L., Steemers, F.J., et al. (2019). The single-cell transcriptional landscape of mammalian organogenesis. Nature 566, 496–502.

15. Chapman, H.A., Li, X., Alexander, J.P., Brumwell, A., Lorizio, W., Tan, K., Sonnenberg, A., Wei, Y., and Vu, T.H. (2011). Integrin alpha6beta4 identifies an adult distal lung epithelial population with regenerative potential in mice. J Clin Invest 121, 2855–2862.

16. Chen, D., Sun, J., Zhu, J., Ding, X., Lan, T., Wang, X., Wu, W., Ou, Z., Zhu, L., Ding, P., et al. (2021). Single cell atlas for 11 non-model mammals, reptiles and birds. Nat Commun 12, 7083.

17. Choi, J., Park, J.E., Tsagkogeorga, G., Yanagita, M., Koo, B.K., Han, N., and Lee, J.H. (2020). Inflammatory Signals Induce AT2 Cell-Derived Damage-Associated Transient Progenitors that Mediate Alveolar Regeneration. Cell Stem Cell 27, 366–382 e367.

18. Danto, S.I., Shannon, J.M., Borok, Z., Zabski, S.M., and Crandall, E.D. (1995). Reversible transdifferentiation of alveolar epithelial cells. Am J Respir Cell Mol Biol 12, 497–502.

19. Demaio, L., Tseng, W., Balverde, Z., Alvarez, J.R., Kim, K.J., Kelley, D.G., Senior, R.M., Crandall, E.D., and Borok, Z. (2009). Characterization of mouse alveolar epithelial cell monolayers. Am J Physiol Lung Cell Mol Physiol 296, L1051–1058.

20. Dobin, A., Davis, C.A., Schlesinger, F., Drenkow, J., Zaleski, C., Jha, S., Batut, P., Chaisson, M., and Gingeras, T.R. (2013). STAR: ultrafast universal RNA-seq aligner. Bioinformatics 29, 15–21.

21. Epshtein, Y., Mathew, B., Chen, W., and Jacobson, J.R. (2023). UCHL1 Regulates Radiation Lung Injury via Sphingosine Kinase-1. Cells 12.

22. Germain, P.L., Lun, A., Garcia Meixide, C., Macnair, W., and Robinson, M.D. (2021). Doublet identification in single-cell sequencing data using scDblFinder. F1000Res 10, 979.

23. Griffiths, J.A., Richard, A.C., Bach, K., Lun, A.T.L., and Marioni, J.C. (2018). Detection and removal of barcode swapping in single-cell RNA-seq data. Nat Commun 9, 2667.

24. Habermann, A.C., Gutierrez, A.J., Bui, L.T., Yahn, S.L., Winters, N.I., Calvi, C.L., Peter, L., Chung, M.I., Taylor, C.J., Jetter, C., et al. (2020). Single-cell RNA sequencing reveals profibrotic roles of distinct epithelial and mesenchymal lineages in pulmonary fibrosis. Sci Adv 6, eaba1972.

25. Hafemeister, C., and Satija, R. (2019). Normalization and variance stabilization of single-cell RNA-seq data using regularized negative binomial regression. Genome Biol 20, 296.

26. Huang, C., Santofimia-Castano, P., and Iovanna, J. (2021). NUPR1: A Critical Regulator of the Antioxidant System. Cancers (Basel) 13.

27. Jaeger, B., Schupp, J.C., Plappert, L., Terwolbeck, O., Artysh, N., Kayser, G., Engelhard, P., Adams, T.S., Zweigerdt, R., Kempf, H., et al. (2022). Airway basal cells show a dedifferentiated KRT17(high)Phenotype and promote fibrosis in idiopathic pulmonary fibrosis. Nat Commun 13, 5637.

28. Jansing, N.L., McClendon, J., Henson, P.M., Tuder, R.M., Hyde, D.M., and Zemans, R.L. (2017). Unbiased Quantitation of Alveolar Type II to Alveolar Type I Cell Transdifferentiation during Repair after Lung Injury in Mice. Am J Respir Cell Mol Biol 57, 519–526.

29. Jiang, P., Gil de Rubio, R., Hrycaj, S.M., Gurczynski, S.J., Riemondy, K.A., Moore, B.B., Omary, M.B., Ridge, K.M., and Zemans, R.L. (2020). Ineffectual Type 2-to-Type 1 Alveolar Epithelial Cell Differentiation in Idiopathic Pulmonary Fibrosis: Persistence of the KRT8(hi) Transitional State. Am J Respir Crit Care Med 201, 1443–1447.

30. Kathiriya, J.J., Wang, C., Zhou, M., Brumwell, A., Cassandras, M., Le Saux, C.J., Cohen, M., Alysandratos, K.D., Wang, B., Wolters, P., et al. (2022). Human alveolar type 2 epithelium transdifferentiates into metaplastic KRT5(+) basal cells. Nat Cell Biol 24, 10–23.

31. Kobayashi, Y., Tata, A., Konkimalla, A., Katsura, H., Lee, R.F., Ou, J., Banovich, N.E., Kropski, J.A., and Tata, P.R. (2020). Persistence of a regeneration-associated, transitional alveolar epithelial cell state in pulmonary fibrosis. Nat Cell Biol 22, 934–946.

32. La Manno, G., Soldatov, R., Zeisel, A., Braun, E., Hochgerner, H., Petukhov, V., Lidschreiber, K., Kastriti, M.E., Lonnerberg, P., Furlan, A., et al. (2018). RNA velocity of single cells. Nature 560, 494–498.

33. LaFemina, M.J., Sutherland, K.M., Bentley, T., Gonzales, L.W., Allen, L., Chapin, C.J., Rokkam, D., Sweerus, K.A., Dobbs, L.G., Ballard, P.L., et al. (2014). Claudin-18 deficiency results in alveolar barrier dysfunction and impaired alveologenesis in mice. Am J Respir Cell Mol Biol 51, 550–558.

34. Lehmann, M., Hu, Q., Hu, Y., Hafner, K., Costa, R., van den Berg, A., and Konigshoff, M. (2020). Chronic WNT/beta-catenin signaling induces cellular senescence in lung epithelial cells. Cell Signal 70, 109588.

35. Lewinska, A., Adamczyk-Grochala, J., Deregowska, A., and Wnuk, M. (2017). Sulforaphane-Induced Cell Cycle Arrest and Senescence are accompanied by DNA Hypomethylation and Changes in microRNA Profile in Breast Cancer Cells. Theranostics 7, 3461–3477.

36. Li, G., Flodby, P., Luo, J., Kage, H., Sipos, A., Gao, D., Ji, Y., Beard, L.L., Marconett, C.N., DeMaio, L., et al. (2014). Knockout mice reveal key roles for claudin 18 in alveolar barrier properties and fluid homeostasis. Am J Respir Cell Mol Biol 51, 210–222.

37. Limbad, C., Doi, R., McGirr, J., Ciotlos, S., Perez, K., Clayton, Z.S., Daya, R., Seals, D.R., Campisi, J., and Melov, S. (2022). Senolysis induced by 25-hydroxycholesterol targets CRYAB in multiple cell types. iScience 25, 103848.

38. Mayr, C.H., Simon, L.M., Leuschner, G., Ansari, M., Schniering, J., Geyer, P.E., Angelidis, I., Strunz, M., Singh, P., Kneidinger, N., et al. (2021). Integrative analysis of cell state changes in lung fibrosis with peripheral protein biomarkers. EMBO Mol Med 13, e12871.

39. Mijit, M., Caracciolo, V., Melillo, A., Amicarelli, F., and Giordano, A. (2020). Role of p53 in the Regulation of Cellular Senescence. Biomolecules 10.

40. Nishi, Y., Sano, H., Kawashima, T., Okada, T., Kuroda, T., Kikkawa, K., Kawashima, S., Tanabe, M., Goto, T., Matsuzawa, Y., et al. (2007). Role of galectin-3 in human pulmonary fibrosis. Allergol Int 56, 57–65.

41. Oka, T., Remue, E., Meerschaert, K., Vanloo, B., Boucherie, C., Gfeller, D., Bader, G.D., Sidhu, S.S., Vandekerckhove, J., Gettemans, J., et al. (2010). Functional complexes between YAP2 and ZO-2 are PDZ domain-dependent, and regulate YAP2 nuclear localization and signalling. Biochem J 432, 461–472.

42. Radwanska, A., Cottage, C.T., Piras, A., Overed-Sayer, C., Sihlbom, C., Budida, R., Wrench, C., Connor, J., Monkley, S., Hazon, P., et al. (2022). Increased expression and accumulation of GDF15 in IPF extracellular matrix contribute to fibrosis. JCI Insight 7.

43. Raslan, A.A., and Yoon, J.K. (2020). WNT Signaling in Lung Repair and Regeneration. Mol Cells 43, 774–783.

44. Riemondy, K.A., Jansing, N.L., Jiang, P., Redente, E.F., Gillen, A.E., Fu, R., Miller, A.J., Spence, J.R., Gerber, A.N., Hesselberth, J.R., et al. (2019). Single cell RNA sequencing identifies TGFbeta as a key regenerative cue following LPS-induced lung injury. JCI Insight 5.

45. Rouaud, F., Sluysmans, S., Flinois, A., Shah, J., Vasileva, E., and Citi, S. (2020). Scaffolding proteins of vertebrate apical junctions: structure, functions and biophysics. Biochim Biophys Acta Biomembr 1862, 183399.

46. Saul, D., Kosinsky, R.L., Atkinson, E.J., Doolittle, M.L., Zhang, X., LeBrasseur, N.K., Pignolo, R.J., Robbins, P.D., Niedernhofer, L.J., Ikeno, Y., et al. (2022). A new gene set identifies senescent cells and predicts senescence-associated pathways across tissues. Nat Commun 13, 4827.

47. Spadaro, D., Tapia, R., Pulimeno, P., and Citi, S. (2012). The control of gene expression and cell proliferation by the epithelial apical junctional complex. Essays Biochem 53, 83–93.

48. Strunz, M., Simon, L.M., Ansari, M., Kathiriya, J.J., Angelidis, I., Mayr, C.H., Tsidiridis, G., Lange, M., Mattner, L.F., Yee, M., et al. (2020). Alveolar regeneration through a Krt8+ transitional stem cell state that persists in human lung fibrosis. Nat Commun 11, 3559.

49. Stuart, T., Butler, A., Hoffman, P., Hafemeister, C., Papalexi, E., Mauck, W.M., 3rd, Hao, Y., Stoeckius, M., Smibert, P., and Satija, R. (2019). Comprehensive Integration of Single-Cell Data. Cell 177, 1888–1902 e1821.

50. Suarez-Artiles, L., Breiderhoff, T., Girardello, R., Gonschior, H., Rodius, S., Lesur, A., Reimer, U., Ramberger, E., Perez-Hernandez, D., Muller, D., et al. (2022). Pan-claudin family interactome analysis reveals shared and specific interactions. Cell Rep 41, 111588.

51. Tam, A.Y.Y., Horwell, A.L., Trinder, S.L., Khan, K., Xu, S., Ong, V., Denton, C.P., Norman, J.T., Holmes, A.M., Bou-Gharios, G., et al. (2021). Selective deletion of connective tissue growth factor attenuates experimentally-induced pulmonary fibrosis and pulmonary arterial hypertension. Int J Biochem Cell Biol 134, 105961.

52. Tamura, A., and Tsukita, S. (2014). Paracellular barrier and channel functions of TJ claudins in organizing biological systems: advances in the field of barriology revealed in knockout mice. Semin Cell Dev Biol 36, 177–185.

53. Ting, C., Aspal, M., Vaishampayan, N., Huang, S.K., Riemondy, K.A., Wang, F., Farver, C., and Zemans, R.L. (2022). Fatal COVID-19 and Non-COVID-19 Acute Respiratory Distress Syndrome Is Associated with Incomplete Alveolar Type 1 Epithelial Cell Differentiation from the Transitional State without Fibrosis. Am J Pathol 192, 454–467.

54. Toth, A., Kannan, P., Snowball, J., Kofron, M., Wayman, J.A., Bridges, J.P., Miraldi, E.R., Swarr, D., and Zacharias, W.J. (2023). Alveolar epithelial progenitor cells require Nkx2-1 to maintain progenitor-specific epigenomic state during lung homeostasis and regeneration. Nat Commun 14, 8452.

55. Trapnell, C., Cacchiarelli, D., Grimsby, J., Pokharel, P., Li, S., Morse, M., Lennon, N.J., Livak, K.J., Mikkelsen, T.S., and Rinn, J.L. (2014). The dynamics and regulators of cell fate decisions are revealed by pseudotemporal ordering of single cells. Nat Biotechnol 32, 381–386.

56. Tsukita, S., and Furuse, M. (2000). The structure and function of claudins, cell adhesion molecules at tight junctions. Ann N Y Acad Sci 915, 129–135.

57. Van Itallie, C.M., and Anderson, J.M. (2013). Claudin interactions in and out of the tight junction. Tissue Barriers 1, e25247.

58. Wabik, A., and Jones, P.H. (2015). Switching roles: the functional plasticity of adult tissue stem cells. EMBO J 34, 1164–1179.

59. Wang, F., Ting, C., Riemondy, K.A., Douglas, M., Foster, K., Patel, N., Kaku, N., Linsalata, A., Nemzek, J., Varisco, B.M., et al. (2023). Regulation of epithelial transitional states in murine and human pulmonary fibrosis. J Clin Invest 133.

60. Wang, Y., Tang, Z., Huang, H., Li, J., Wang, Z., Yu, Y., Zhang, C., Li, J., Dai, H., Wang, F., et al. (2018). Pulmonary alveolar type I cell population consists of two distinct subtypes that differ in cell fate. Proc Natl Acad Sci U S A 115, 2407–2412.

61. Weiler, P., Lange, M., Klein, M., Pe’er, D., and Theis, F. (2024). CellRank 2: unified fate mapping in multiview single-cell data. Nat Methods 21, 1196–1205.

62. Wolf, F.A., Angerer, P., and Theis, F.J. (2018). SCANPY: large-scale single-cell gene expression data analysis. Genome Biol 19, 15.

63. Xu, Y., Wang, Y., Besnard, V., Ikegami, M., Wert, S.E., Heffner, C., Murray, S.A., Donahue, L.R., and Whitsett, J.A. (2012). Transcriptional programs controlling perinatal lung maturation. PLoS One 7, e37046.

64. Yeo, H.J., Ha, M., Shin, D.H., Lee, H.R., Kim, Y.H., and Cho, W.H. (2024). Development of a Novel Biomarker for the Progression of Idiopathic Pulmonary Fibrosis. Int J Mol Sci 25.

65. Zhang, H.Y., Gharaee-Kermani, M., Zhang, K., Karmiol, S., and Phan, S.H. (1996). Lung fibroblast alpha-smooth muscle actin expression and contractile phenotype in bleomycin-induced pulmonary fibrosis. Am J Pathol 148, 527–537.

66. Zhao, B., Li, L., Lu, Q., Wang, L.H., Liu, C.Y., Lei, Q., and Guan, K.L. (2011). Angiomotin is a novel Hippo pathway component that inhibits YAP oncoprotein. Genes Dev 25, 51–63.

67. Zhao, L., Yee, M., and O’Reilly, M.A. (2013). Transdifferentiation of alveolar epithelial type II to type I cells is controlled by opposing TGF-beta and BMP signaling. Am J Physiol Lung Cell Mol Physiol 305, L409–418.

68. Zhou, B., Flodby, P., Luo, J., Castillo, D.R., Liu, Y., Yu, F.X., McConnell, A., Varghese, B., Li, G., Chimge, N.O., et al. (2018). Claudin-18-mediated YAP activity regulates lung stem and progenitor cell homeostasis and tumorigenesis. J Clin Invest 128, 970–984.

69. Zhou, B., Stueve, T.R., Mihalakakos, E.A., Miao, L., Mullen, D., Wang, Y., Liu, Y., Luo, J., Tran, E., Siegmund, K.D., et al. (2021). Comprehensive epigenomic profiling of human alveolar epithelial differentiation identifies key epigenetic states and transcription factor co-regulatory networks for maintenance of distal lung identity. BMC Genomics 22, 906.

