## Supplemental for "Loss of tight junction protein claudin 18 uncovers alveolar epithelial stem cell plasticity and emergence of non-fibrogenic transitional progenitors"

**Supplemental Figure 1 – Principal component selection and marker gene expression in AECs.** A) Elbow plot of principal components (PC) showing that the variance in gene expression begins to plateau around PC30, which was selected as the cutoff for downstream analyses. B) Heatmap of expression of cell population markers in AECs.

**Supplemental Figure 2 – KRT8 is expressed in adult *Cldn18* KO lungs in the absence of fibrosis.** IF (A) and quantitation (B) for KRT8 in *Cldn18* KO and WT lungs of adult over time. Scale bars = 50 µm; N = 3-5. * = $p < 0.05$, **= $p < 0.01$, **** = $p < 0.0001$. Two-way ANOVA. C) Low magnification view of Figure 1 C: Sirius red staining showing signal around the airways in both WT and *Cldn18* KO lungs as a positive control.

**Supplemental Figure 3 - $KRT8^+$ and $KRT19^+$ cells express markers of AT1 and AT2 cells.** A) IF showing lower magnification of images in Figure 1 E-F. Dashed white square = magnified areas in Figure 1 E-F. Scale bars = 50 µm; N = 6. B-C) Quantitation of A: $AQP5^+KRT8^+$ and $AQP5^+KRT19^+$ as a percentage of $AQP5^+$ cells in WT and *Cldn18* KO adult lung (B); n = 4-6; ** = $p < 0.01$. Unpaired t-test. $SFTPC^+$ $KRT8^+$ and $SFTPC^+$ $KRT19^+$ as a percentage of $SFTPC^+$ cells in WT and *Cldn18* KO adult lungs (C); N= 4-7; NS = not significant. Unpaired t-test.

**Supplemental Figure 4 – *Cldn18* KO AECs are susceptible to bleomycin.** Western blot (WB) (A) and quantitation (B) of P-histone-H2AX expression in AEC in culture treated with bleomycin at day 3-4 (n=3) and day 6-7 (n=2). NS = not significant; **= $p \leq 0.01$, *** $p \leq 0.001$, **** = $p \leq 0.0001$. Two-way ANOVA.

**Supplemental Figure 5 – Single cell RNA-seq captured the full set of lung epithelial populations.** Combined UMAP of WT and *Cldn18* KO $EPCAM^+$ cells.

**Supplemental Figure 6 – RATPs differ from DATPs**. Violinplot for markers in Figure 4 B show significant differences between DATPs and RATPs for markers of AT2 (A), transitional (B) and

AT1 (C) cell markers, in support of the dotplot in Figure 4 B. * = p < 0.05, **= p < 0.01, *** = p < 0.001, **** = p < 0.0001. Unpaired t-test.

**Supplemental Figure 7 – RATPs in other published datasets.** UMAP of integrated analysis of our dataset with LPS (Riemondy et al., 2019), organoids (Kobayashi et al., 2020), and long-term bleomycin (Strunz et al., 2020) datasets.

**Supplemental Figure 8 - Downregulation of *Nupr1* in RATPs vs DATPs.** Gene expression and prediction of its downregulation by upstream analysis (Ingenuity Pathway Analysis) in RATPs vs DATPs. Green: downregulated; red: upregulated.

**Supplemental Figure 9 - ScRNA-seq biological replicates** – A) UMAPs show biological replicates for WT and *Cldn18* KO samples following bleomycin. B) Violin plots showing expression of AEC markers across biological replicates.

**Supplemental Figure 10 – Restoration of homeostatic gene expression in AEC populations of *Cldn18* KO by D7 following bleomycin**. Number of genes altered in each cell population at D3 vs D0, D7 vs D3 and D7 vs D0 in WT (circles) and *Cldn18* KO (triangles) samples.

**Supplemental Figure 11 – Expanded inflamed AT2 cell population in both WT and *Cldn18* KO lungs following bleomycin injury.** A) Percentage of inflamed AT2 cells over time. B) UMAP showing expression of inflammatory markers in WT and *Cldn18* KO mice over time.

**Supplementary Figure 12 – RATP1s and RATP2s are distinct from AT1 and AT2 cell populations**. A) Violin plots for AT2 cell markers. B) Violin plots for AT1 cell markers

**Supplementary Figure 13 - Accelerated AEC differentiation in the lungs of *Cldn18* KO compared to WT** A) Violin plot of diffusion pseudotime (DPT) of WT and *Cldn18* KO scRNA-seq datasets. B) RNA velocity of WT and *Cldn18* KO scRNA-seq datasets. C) *tdTomato* signal for the *Sftpc*-lineage trace at D0, D3 and D7 following injury in AT1 cells.

#### Supplemental Figure 1

**A**

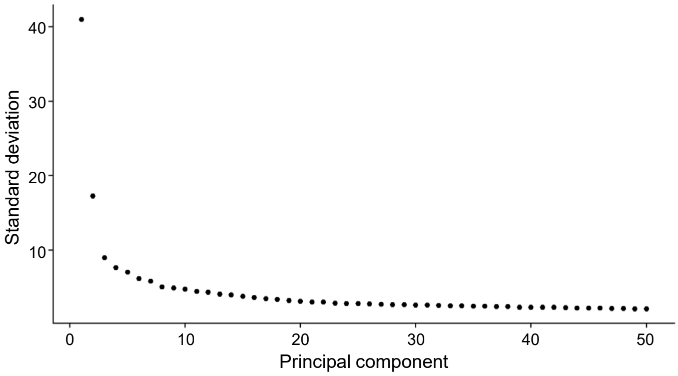


**B**

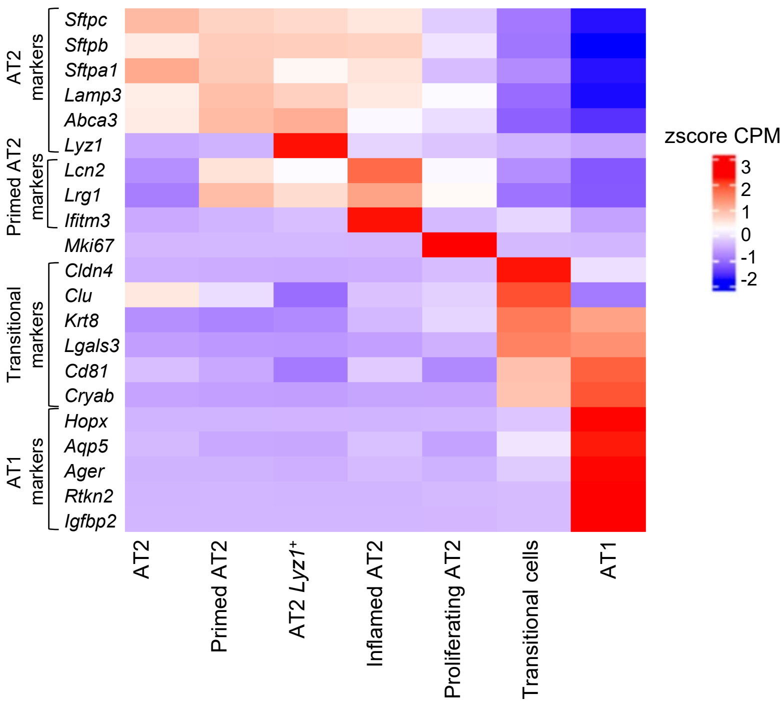

### Supplemental Figure 2

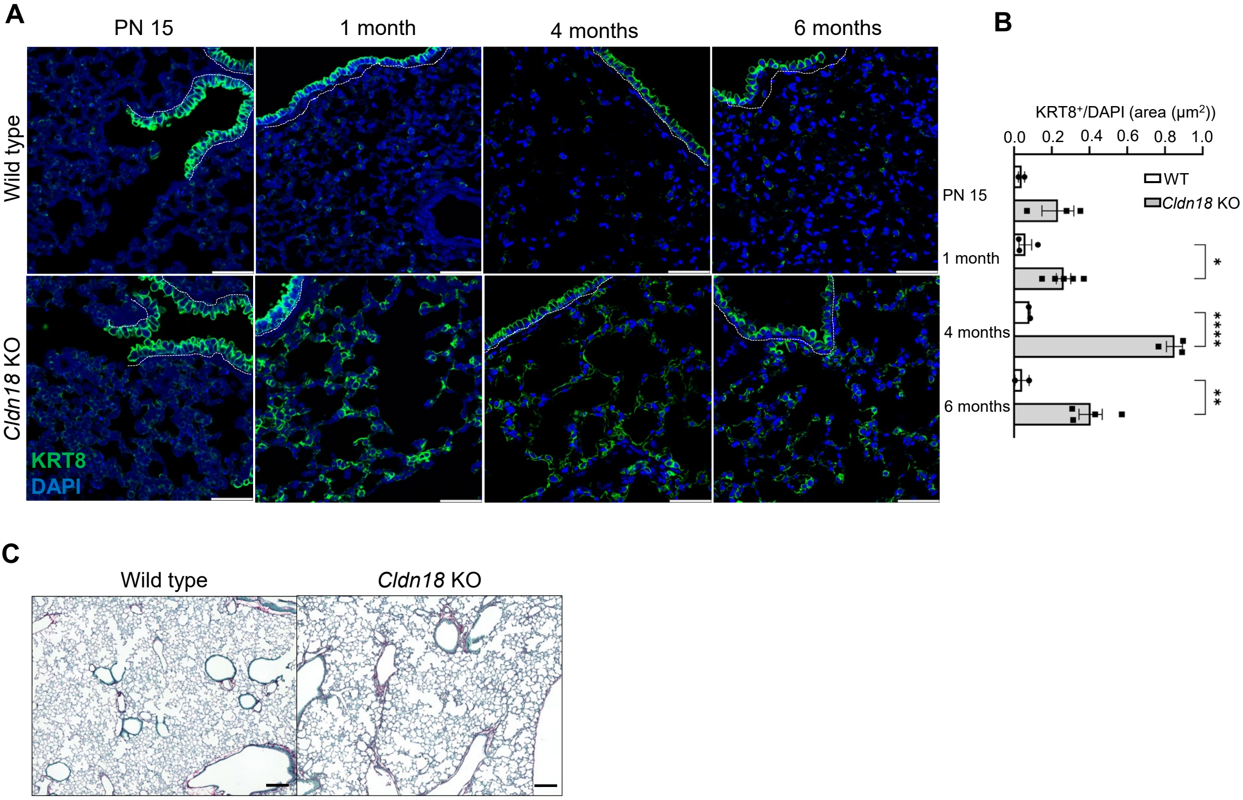

**Supplemental Figure 3**

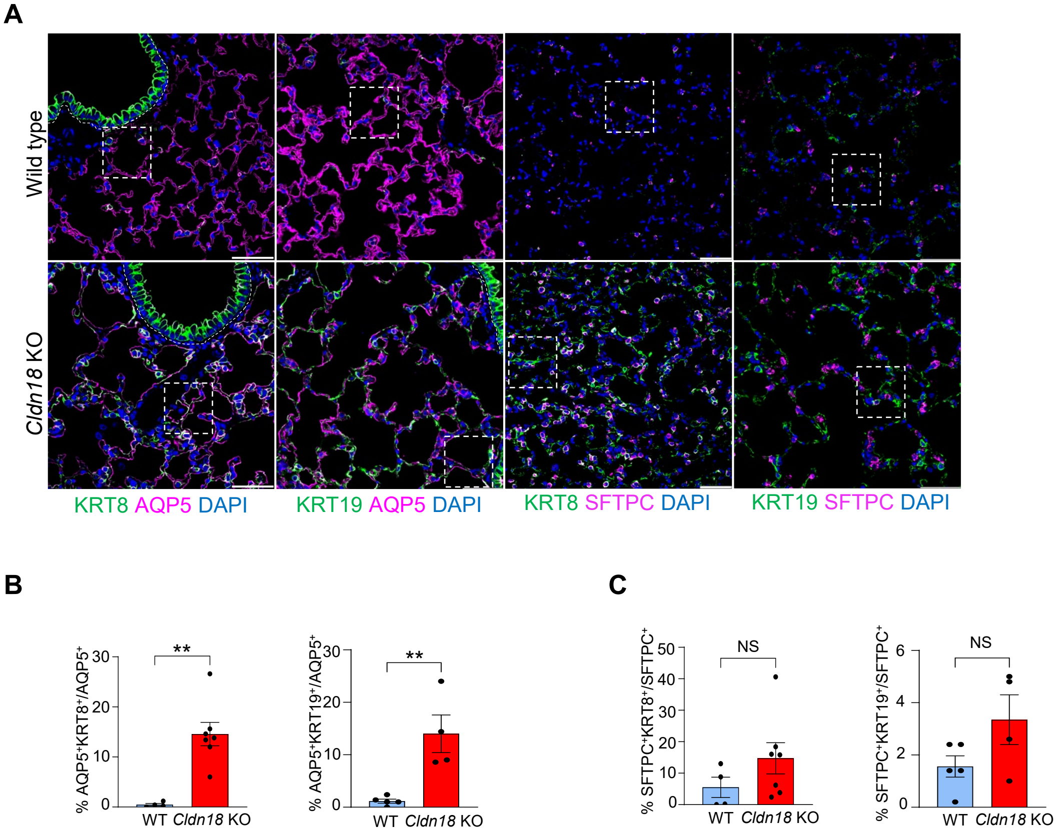

#### Supplemental Figure 4

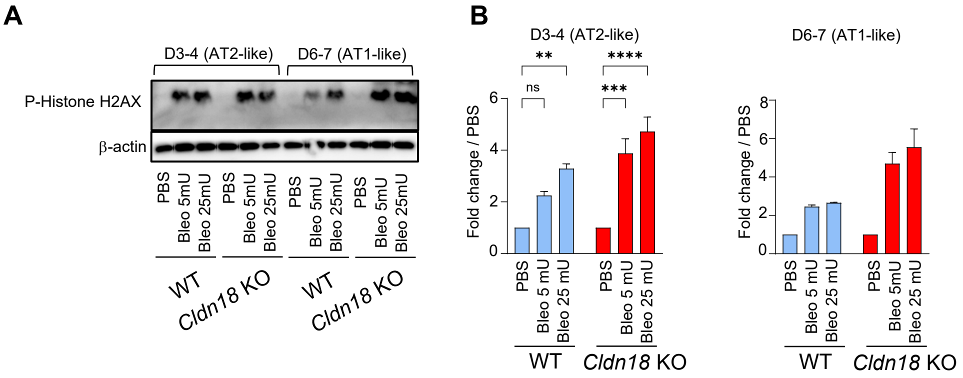

#### Supplemental Figure 5

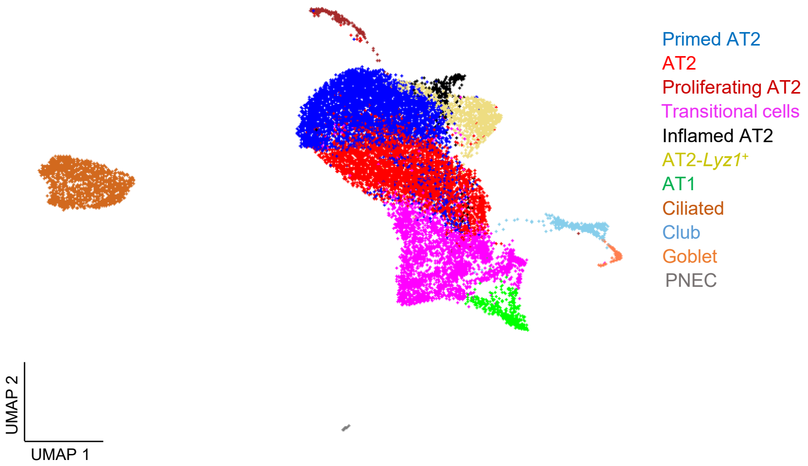

### Supplemental Figure 6

**A**

AT2 markers

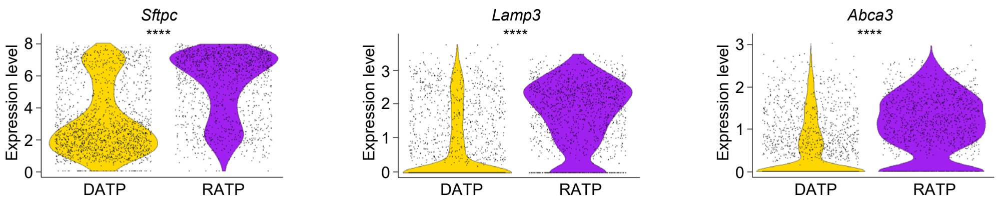


**B**

Transitional markers

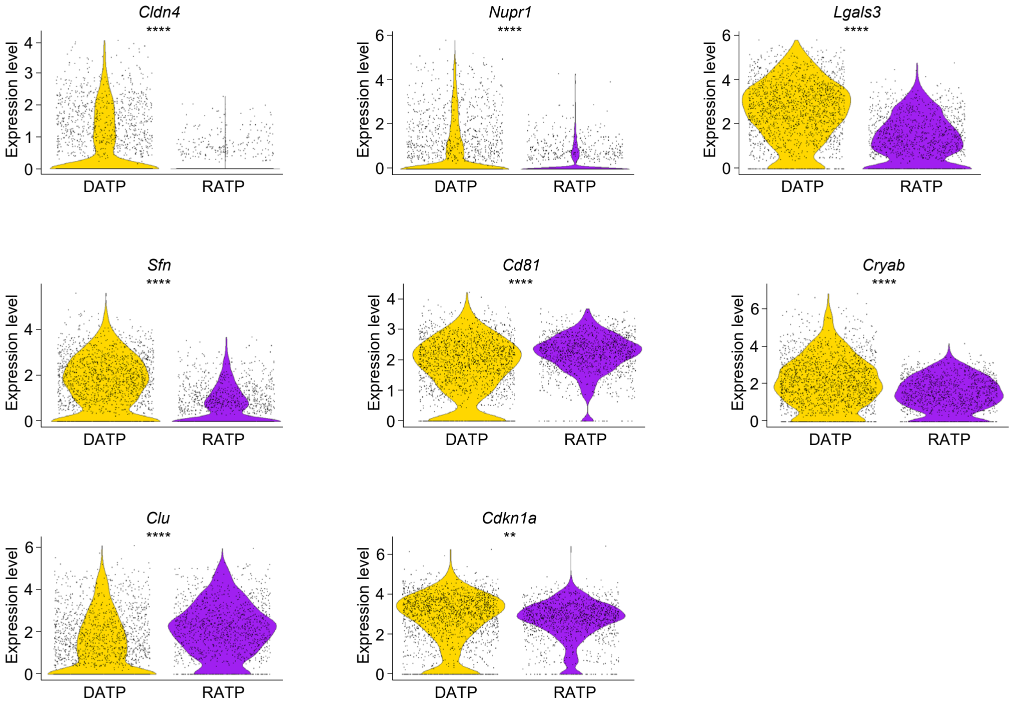


**C**

AT1 markers

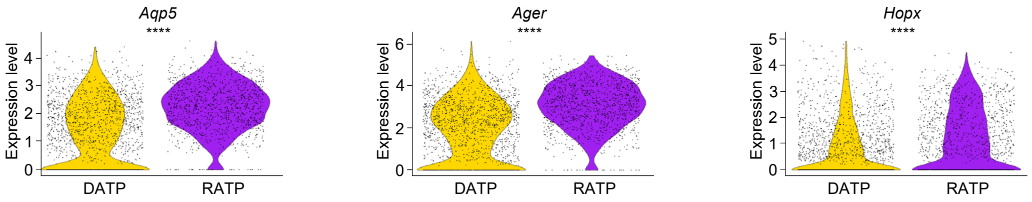

#### Supplemental Figure 7

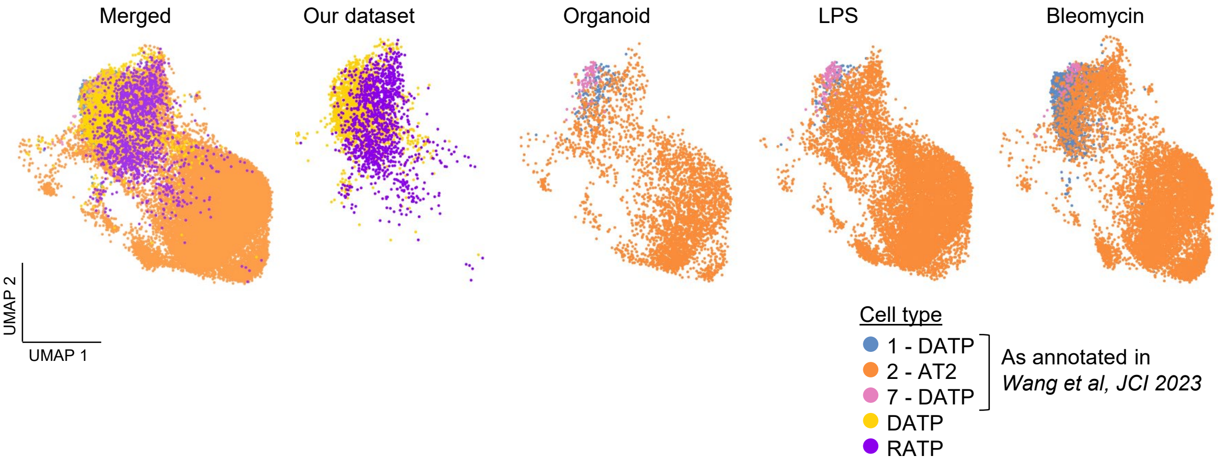

#### Supplemental Figure 8

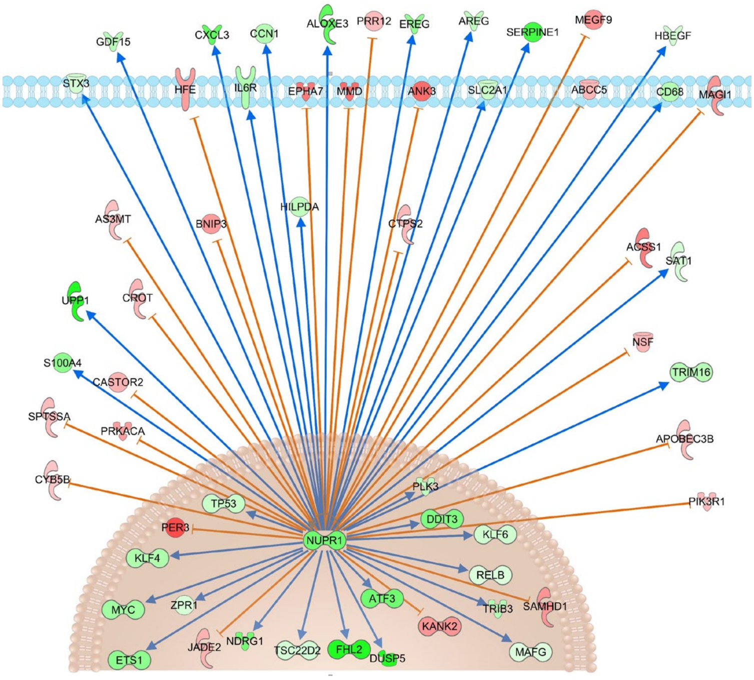

**Supplemental Figure 9**

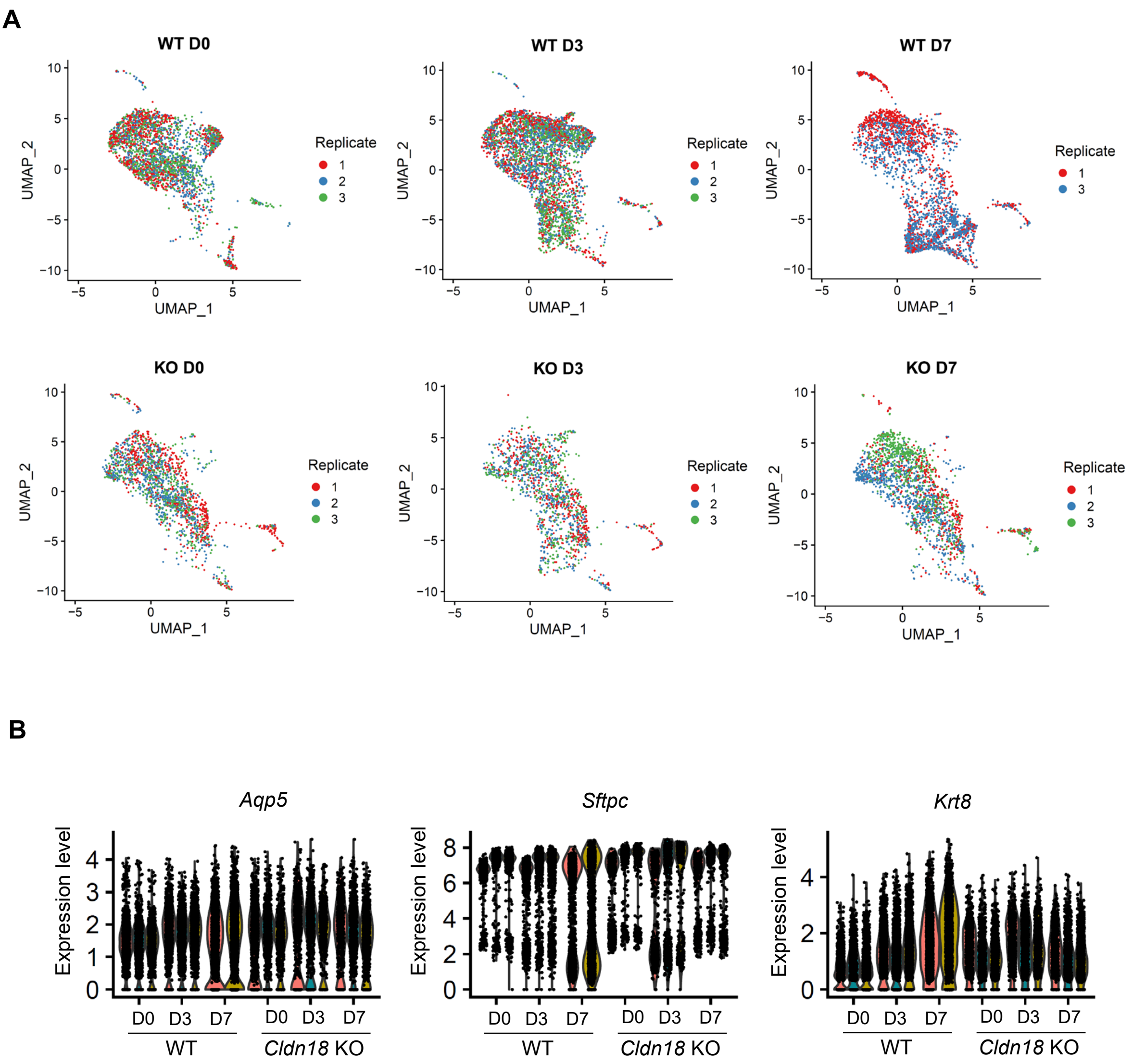

#### Supplemental Figure 10

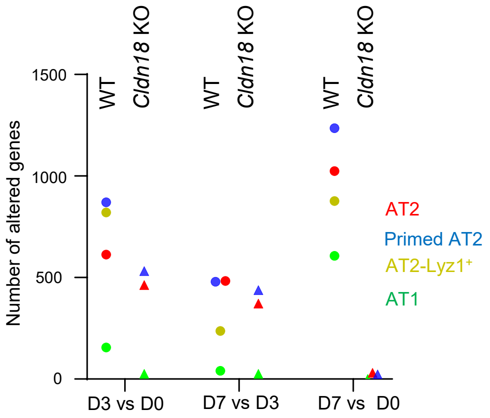

**Supplemental Figure 11**

A

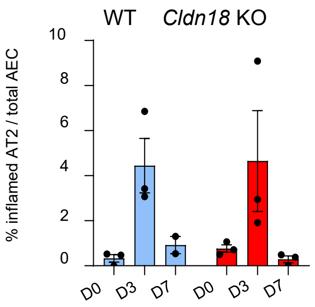


B

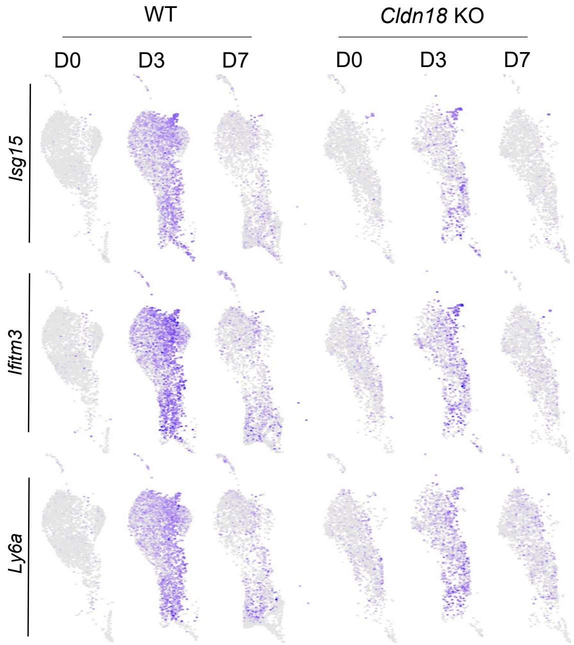

**Supplementary Figure 12**

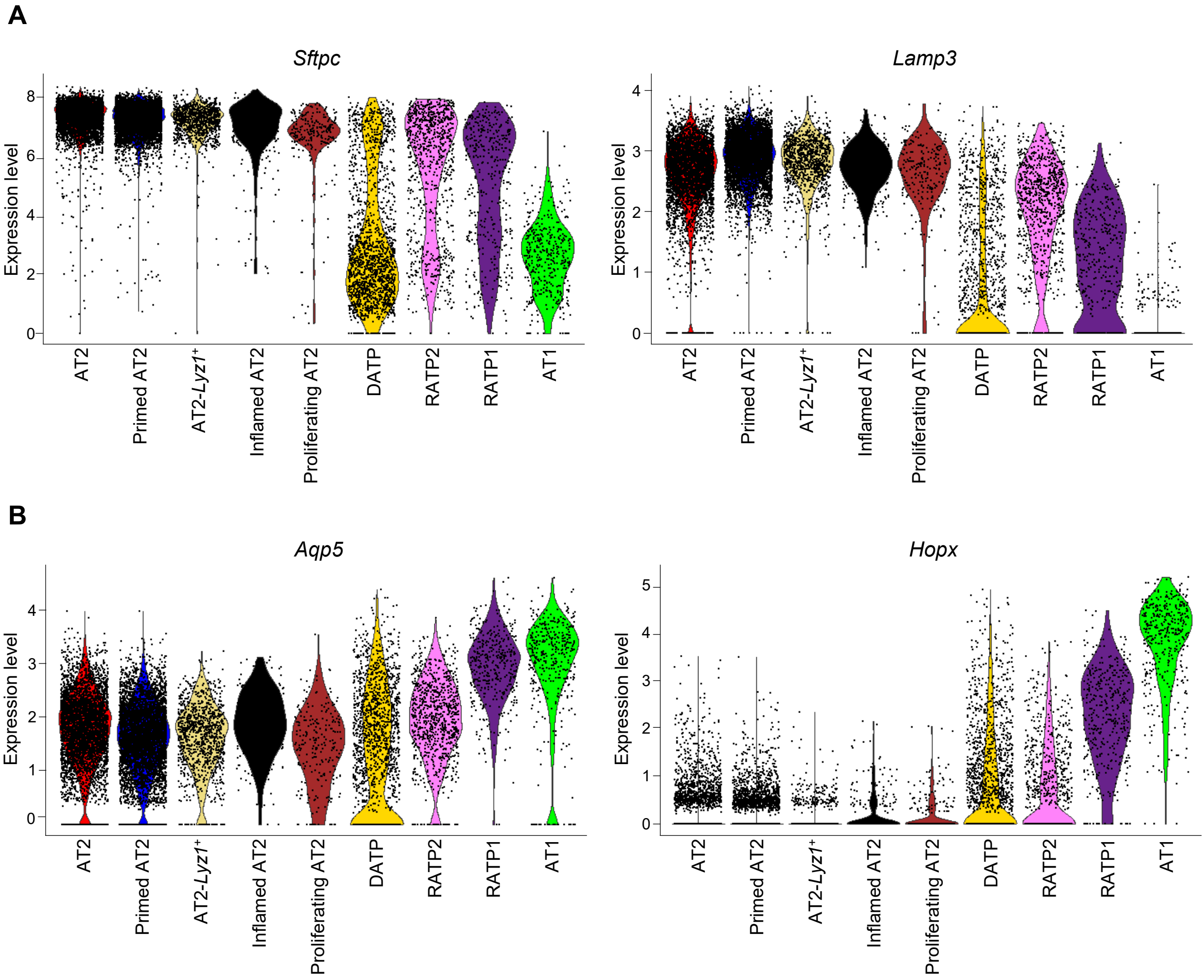

#### Supplementary Figure 13

A

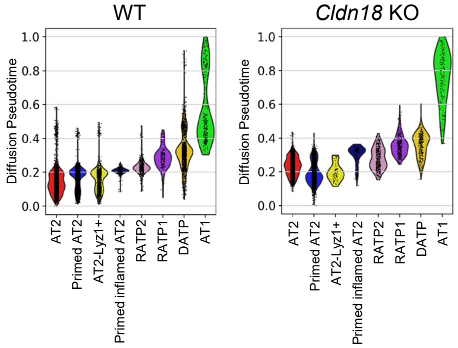


B

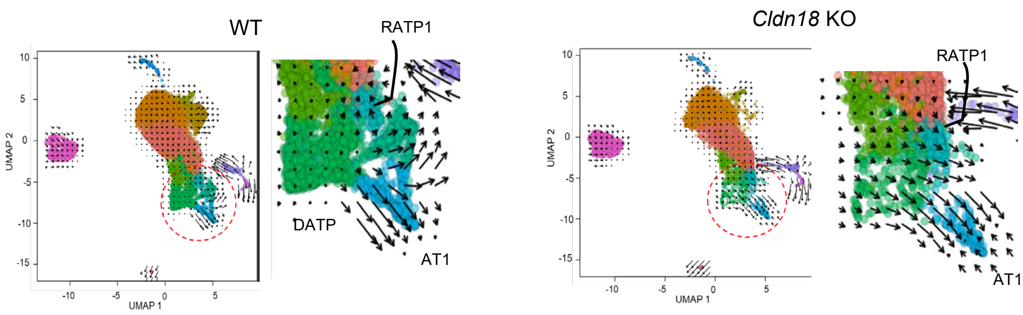


C

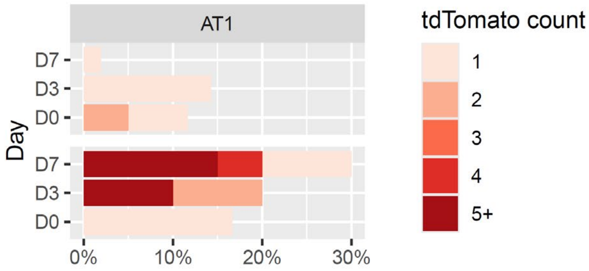
